# Pattern-recognition receptors are required for NLR-mediated plant immunity

**DOI:** 10.1101/2020.04.10.031294

**Authors:** Minhang Yuan, Zeyu Jiang, Guozhi Bi, Kinya Nomura, Menghui Liu, Sheng Yang He, Jian-Min Zhou, Xiu-Fang Xin

## Abstract

The plant immune system is fundamental to plant survival in natural ecosystems and productivity in crop fields. Substantial evidence supports the prevailing notion that plants possess a two-tiered innate immune system, called pattern-triggered immunity (PTI) and effector-triggered immunity (ETI). PTI is triggered by microbial patterns via cell surface-localized pattern-recognition receptors (PRRs), whereas ETI is activated by pathogen effector proteins via mostly intracellularly-localized receptors called nucleotide-binding, leucine-rich repeat proteins (NLRs)^1-4^. PTI and ETI are initiated by dist 30 inct activation mechanisms and are considered to act independently and have evolved sequentially^5,6^. Here we show that, contrary to the perception of PTI and ETI being separate immune signaling pathways, Arabidopsis PRR/co-receptor mutants, *fls2/efr/cerk1* and *bak1/bkk1/cerk1* triple mutants, are greatly impaired in ETI responses when challenged with incompatible *Pseudomonas syrinage* bacteria. We further show that the NADPH oxidase (RBOHD)-mediated production of reactive oxygen species (ROS) is a critical early signaling event connecting PRR and NLR cascades and that PRR-mediated phosphorylation of RBOHD is necessary for full activation of RBOHD during ETI. Furthermore, NLR signaling rapidly augments the transcript and protein levels of key PTI components at an early stage and in a salicylic acid-independent manner. Our study supports an alternative model in which PTI is in fact an indispensable component of ETI during bacterial infection, implying that ETI halts pathogen infection, in part, by directly co-opting the anti-pathogen mechanisms proposed for PTI. This alternative model conceptually unites two major immune signaling pathways in the plant kingdom and mechanistically explains the long-observed similarities in downstream defense outputs between PTI and ETI.

## Main

Signaling initiated by PRRs and NLRs lead to largely overlapping cellular features, but the mechanism(s) by which this occurs and the nature of the signal collaboration between cell surface and intracellular perception systems has remained undiscovered. PRRs are cell surface-localized receptor-like kinases/proteins (RLKs/RLPs) with extracellular ligand-binding domain to sense conserved molecular patterns, ranging from bacterial flagellin to fungal chitin molecules from both pathogenic and nonpathogenic microbes. NLRs, on the other hand, are intracellular proteins that sense pathogen-derived effector proteins inside the plant cell and can be further classified into the coiled coil (CC)-type, Toll/interleukin-1 receptor (TIR)-type, or RPW8 (CC_R_)-type, depending on their N-terminal domain^7^. However, PRR- and NLR-mediated signaling pathways result in many similar downstream immune outputs, including defense gene expression, production of ROS and callose deposition at the plant cell wall^8,9^. The underlying reason is not clear and mechanistic relationship between the two immune pathways remains largely enigmatic. Notably, while many PRR signaling components have been identified and anti-pathogen mechanisms described, the downstream signaling events in ETI and how ETI halts pathogen growth still remain poorly understood, despite recent breakthroughs in the understanding of NLR protein structures and activities^10-13^.

### Requirement of PRR/co-receptors for ETI

Using *Arabidopsis thaliana-Pseudomonas syringae* pathosystem, we accidentally discovered a striking and unexpected role of PRR/co-receptors in ETI. Specifically, an “avirulent”, ETI-eliciting bacterial strain, *P. syringae* pv. *tomato* (*Pst*) DC3000(*avrRpt2*), which activates RPS2 (Resistance to *P. syringae* 2)-dependent ETI in wild-type Col-0 plants^14,15^, failed to elicit effective ETI in two separate PRR/co-receptor Arabidopsis mutants, *fls2/efr/cerk1* (*fec*) and *bak1/bkk1/cerk1* (*bbc*) mutants, which are mutated in major PRR/co-receptors recognizing bacteria-associated molecular patterns^16^. As shown in Fig. 1a, the *fec* and *bbc* mutants did not mount an effective ETI against *Pst* DC3000(*avrRpt2*). To determine whether a requirement of PRR/co-receptors for ETI is specific to *Pst* DC3000(*avrRpt2*) or is a more general phenomenon, we tested two other ETI-triggering “avirulent” effectors, AvrPphB and AvrRps4, which are recognized by RPS5^17^ and RPS4^18^, respectively, in Arabidopsis Col-0 accession. We found that the compromised ETI phenotype in *fec* and *bbc* mutants held true for both AvrPphB and AvrRps4 (Extended Data Fig. 1), suggesting a potentially broad role of PRR/co-receptors in ETI pathways. We subsequently focused on AvrRpt2-triggered ETI for in-depth characterization. Hypersensitive response (HR), manifested by fast cell death under high bacterial inoculum, is a hallmark of ETI. We tested HR phenotype in *fec* and *bbc* mutants in response to *Pst* DC3000(*avrRpt2*) and found that, even though HR cell death eventually occurs in these mutants, the rate was delayed, as shown by the compromised tissue collapse 7h after bacteria infiltration (Fig. 1b).

**Fig.1.**
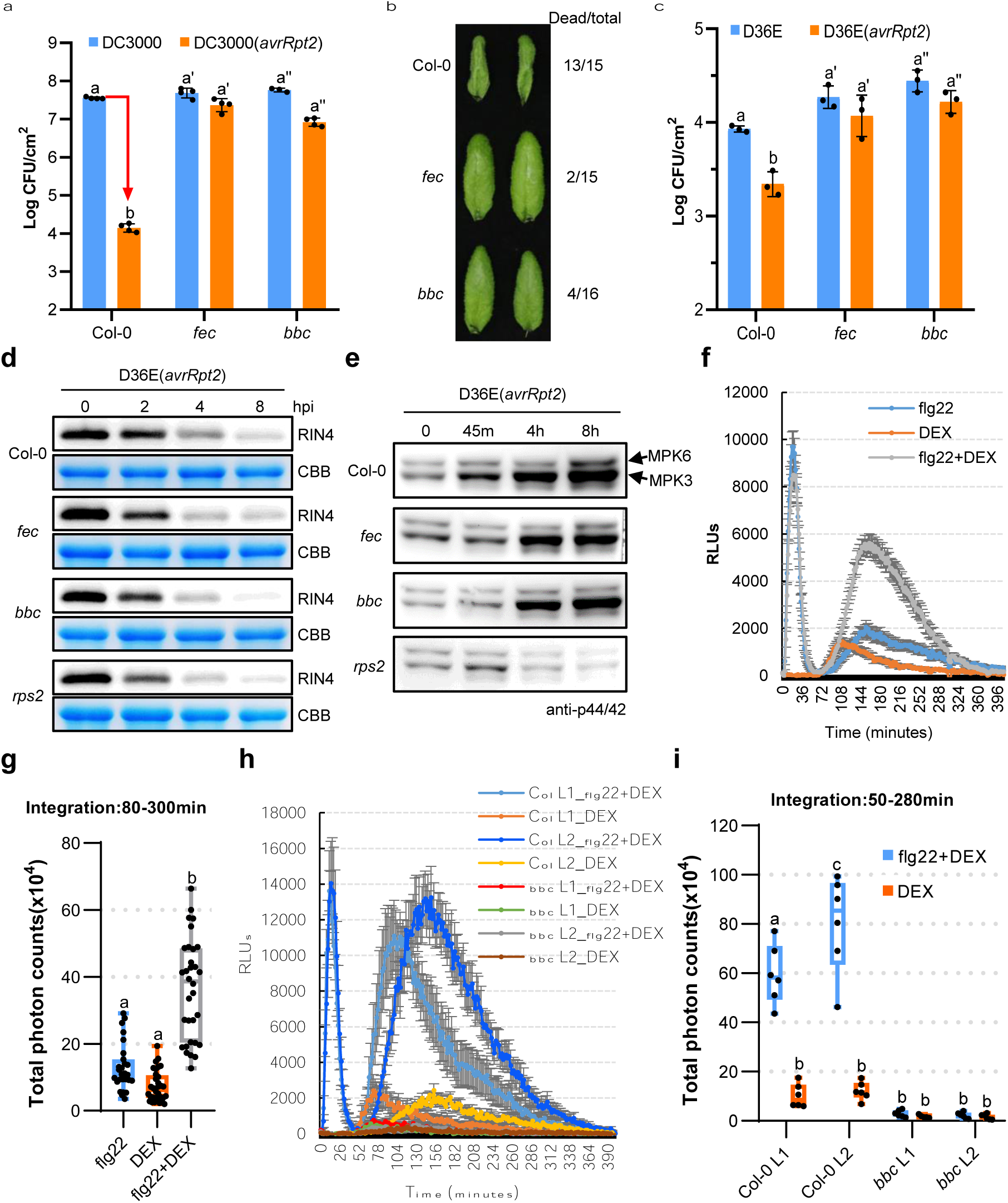
PTI-associated PRR/co-receptors are required for ETI responses and resistance. **a**, Two independent triple PRR/co-receptor mutant plants were hyper-susceptible to *Pst* DC3000 (*avrRpt2*) infection. Bacteria were infiltrated into Arabidopsis leaves at OD_600_=0.002 and bacterial populations inside leaves were determined 3 days post infection. **b**, HR cell death (indicated by leaf collapse in Col-0) was compromised in PRR/co-receptor mutants. *Pst* DC3000 (*avrRpt2*) bacteria were infiltrated into Arabidopsis leaves at OD_600_=0.2 and pictures were taken ∼7 h after infiltration. Numbers of dead and total infiltrated leaves were counted. **c**, The triple PRR/co-receptor mutant plants were hyper-susceptible to D36E(*avrRpt2*) infection. Bacteria were infiltrated into Arabidopsis leaves at OD_600_=0.004 and bacterial populations inside leaves were determined 4 days post infection. Different letters in **a**, and **c**, indicate statistically significant differences in bacterial population, as determined by two-way ANOVA (mean ± s.d.; *n* ≥ 3; *p* < 0.05). **d, e**, RIN4 cleavage **d** and MPK3/6 phosphorylation **e** in Col-0 and the PRR/co-receptor mutants after D36E(*avrRpt2*) inoculation. CBB, Coomassie Brilliant Blue staining. An equal amount of total protein was loaded in each lane. **f, g**, ROS burst detected by luminol-HRP approach in the DEX-inducible *avrRpt2* transgenic plant after treatment of 100nM flg22, 5μM DEX, or both. RLUs, relative luminescence units. Total photon counts are calculated from **f** (*n* ≥ 27). Different letters indicate statistically significant differences as analyzed by one-way ANOVA (*p* < 0.05). **h, i**, ROS burst in Col-0/*DEX::avrRpt2* and *bbc*/*DEX::avrRpt2* plants after treatment of 5μM DEX, or 100nM flg22+5μM DEX. Total photon counts are calculated from **h** (*n* ≥ 5). Different letters indicate statistically significant differences as analyzed by two-way ANOVA (*p* < 0.05). Box plots: centre line, median; box limits, lower and upper quartiles; whiskers, highest and lowest data points. Dots indicate individual data points. All experiments were repeated at least three times with similar trends.

For the past several decades, conventional studies of ETI triggered by *Pst* DC3000 carrying “avirulent” effector genes have been performed in the presence of all 36 endogenous effector genes in *Pst* DC3000. Because during pathogen infection, both PTI and ETI are at play and because many of the endogenous effectors in *Pst* DC3000 are linked to interference of PTI and/or ETI via unclear mechanisms, it is not always easy to clearly interpret the relationship between PTI and ETI during infection by avirulent *Pst* DC3000 strains or other wild-type avirulent pathogens^19-21^. We took advantage of the recent availability of *Pst* DC3000 strain D36E^22^, in which all 36 effector genes as well as coronatine biosynthesis genes are deleted and therefore is expected to activate only PTI, and D36E(*avrRpt2*) strain, which delivers only AvrRpt2 and activates both PTI and RPS2-mediated ETI with no interference from any endogenous *Pst* DC3000 effectors. Although D36E is greatly reduced in virulence compared to *Pst* DC3000 (Fig. 1a), we could still observe a robust AvrRpt2-induced ETI in Col-0 plants, with strain D36E(*avrRpt2*) growing significantly less than strain D36E (Fig. 1c). We found that AvrRpt2-triggered ETI was no longer detectable in either *fec* or *bbc* mutant (Fig. 1c), demonstrating that the requirement of PRR/co-receptors for AvrRpt2-triggered ETI was not caused by some hidden interactions between endogenous *Pst* DC3000 effectors/coronatine and Arabidopsis *fec* and *bbc* mutants.

### Requirement of PRR/co-receptors for ROS production in ETI

Previous studies have shown that AvrRpt2 cleaves the plant protein RIN4 (RPM1-interacting protein 4), leading to activation of the NLR protein RPS2^14,15^. To understand why AvrRpt2-mediated ETI is lost in *fec* and *bbc* mutants, we examined the cleavage of the RIN4 protein by AvrRpt2. Results showed that D36E(*avrRpt2*)-induced RIN4 protein depletion was normal in the *fec* and *bbc* mutants compared to that in Col-0 plants (Fig. 1d). Gene expression analysis showed that *RPS2* transcript level is also similar in all genotypes after bacteria inoculation (Extended Data Fig. 2). We then sought to assay downstream signaling events of AvrRpt2-triggered ETI and examined MAPK phosphorylation level. Activation of RPS2 by AvrRpt2 is known to trigger a strong and more sustained MAPK activation, compared to that induced during PTI^23^. As shown in Fig. 1e, we observed strong ETI-associated MPK3/6 phosphorylation (i.e., at 4 or 8 h post inoculation) in Col-0 plants, which however remained intact in *fec* and *bbc* mutants.

Another important immune response associated with both PTI and ETI is production of ROS, including superoxide and hydrogen peroxide (H_2_O_2_), which have been proposed to act as defense molecules that kill pathogens and signaling molecules that further activate immune responses^24^. Using luminol-horseradish peroxidase (HRP)-based method, we examined PTI- and ETI-associated ROS production in transgenic *avrRpt2* plants, in which *avrRpt2* expression is driven by a dexamethasone (DEX)-inducible promoter^25^. In this system, PTI and ETI activation can be initiated separately or in combination using PAMP (e.g., flg22, a 22-aa peptide derived from bacterial flagellin) and DEX treatments in the absence of bacterial infection. As shown in Fig. 1f, flg22 alone triggered a fast and transient ROS burst as reported before^26^, while DEX-induced expression of AvrRpt2 alone triggered only a weak and kinetically slower ROS burst. Interestingly, co-treatment of flg22 and DEX triggered a strong and sustained second-phase ROS burst, peaking at 2h to 3h after treatment, and lasted for several hours (Fig. 1f, g), a profile that bears a striking similarity to previous observations during bacteria-triggered ETI^27,28^. This result raises an intriguing possibility that activation of PTI may be required for the production of a strong and sustained ROS characteristic of ETI. To test this hypothesis, we generated *bbc*/*DEX::avrRpt2* plants by transforming the *DEX::avrRpt2* construct into the *bbc* mutant plant (as well as Col-0 plant as control). Independent lines in which the *avrRpt2* expression level was similar or even higher than that in Col-0/*DEX::avrRpt2* plants were chosen (Extended Data Fig. 3) for further analysis. As shown in Fig. 1h, i, in the *bbc*/*DEX::avrRpt2* plants, not only flg22-induced first-phase ROS is absent, but also the second-phase AvrRpt2-triggered ROS burst is almost completely abolished, clearly demonstrating a requirement of PRR/co-receptors for ETI-associated ROS production.

To examine whether PTI- and ETI-associated ROS bursts are produced at the same or different subcellular compartments, ROS production was monitored with the fluorescent dye H_2_DCFDA, which can cross the plasma membrane of the plant cell and detect both apoplastic and cytosolic ROS^29^. As shown in Fig. 2a, strong fluorescent signal was detected in the apoplastic space of Col-0 leaves 5h post infiltration of D36E(*avrRpt2*). The apoplastic signal was much weaker in the *bbc* mutant plant, which was indistinguishable compared to the *rps2* control plant infiltrated with D36E(*avrRpt2*) or Col-0 plant infiltrated with D36E (Fig. 2a). Two classes of enzymes, the NADPH oxidases such as respiratory burst oxidase homolog D (RBOHD) and peroxidases, have been shown to be involved in generating apoplastic ROS in pathogen-infected plant leaves^30,31^. We therefore investigated which class is involved in the generation of AvrRpt2-triggered ROS by using chemical inhibitors diphenylene iodonium (DPI), which inhibits NADPH oxidases, and salicylhydroxamic acid (SHAM) and sodium azide, which inhibit peroxidase activities^27,32^. As shown in Extended Data Fig. 4a-c, co-treatment of DPI, but not SHAM or sodium azide, with flg22 and DEX almost completely blocked ETI-associated ROS and greatly compromised PTI-associated ROS. Interestingly, when we added these inhibitors at 40 min after flg22+DEX treatment (i.e., after PTI-associated ROS and before the start of ETI-associated ROS), still only DPI, but not SHAM or sodium azide, almost completely blocked ETI-ROS (Fig. 2b), indicating that NADPH oxidases mediate ETI-associated ROS. We further tested whether NADPH oxidase RBOHD, which has been shown to play a prominent role in generating pathogen-induced ROS^30,33,34^, mediates the ETI-associated ROS. As shown in Fig. 2c, D36E(*avrRpt2*)-induced apoplastic ROS, as detected *in planta* by the H_2_DCFDA dye, was completely lost in the *rbohd* plant. Consistently, we detected a much compromised ETI resistance against *Pst* DC3000(*avrRpt2*) in the *rbohd* mutant plant (Fig. 2d, Extended Data Fig. 5b). Notably, *rbohd* mutant plants grew to sizes that were similar to wild-type Col-0 plants under optimized growth conditions and, under these conditions, showed similar or slightly enhanced susceptibility to *Pst* DC3000 (Fig. 2d, Extended Data Fig. 5a, b). Altogether, our results demonstrate a critical role of RBOHD in ETI and suggests RBOHD as a key molecular node connecting PTI and ETI.

**Fig.2.**
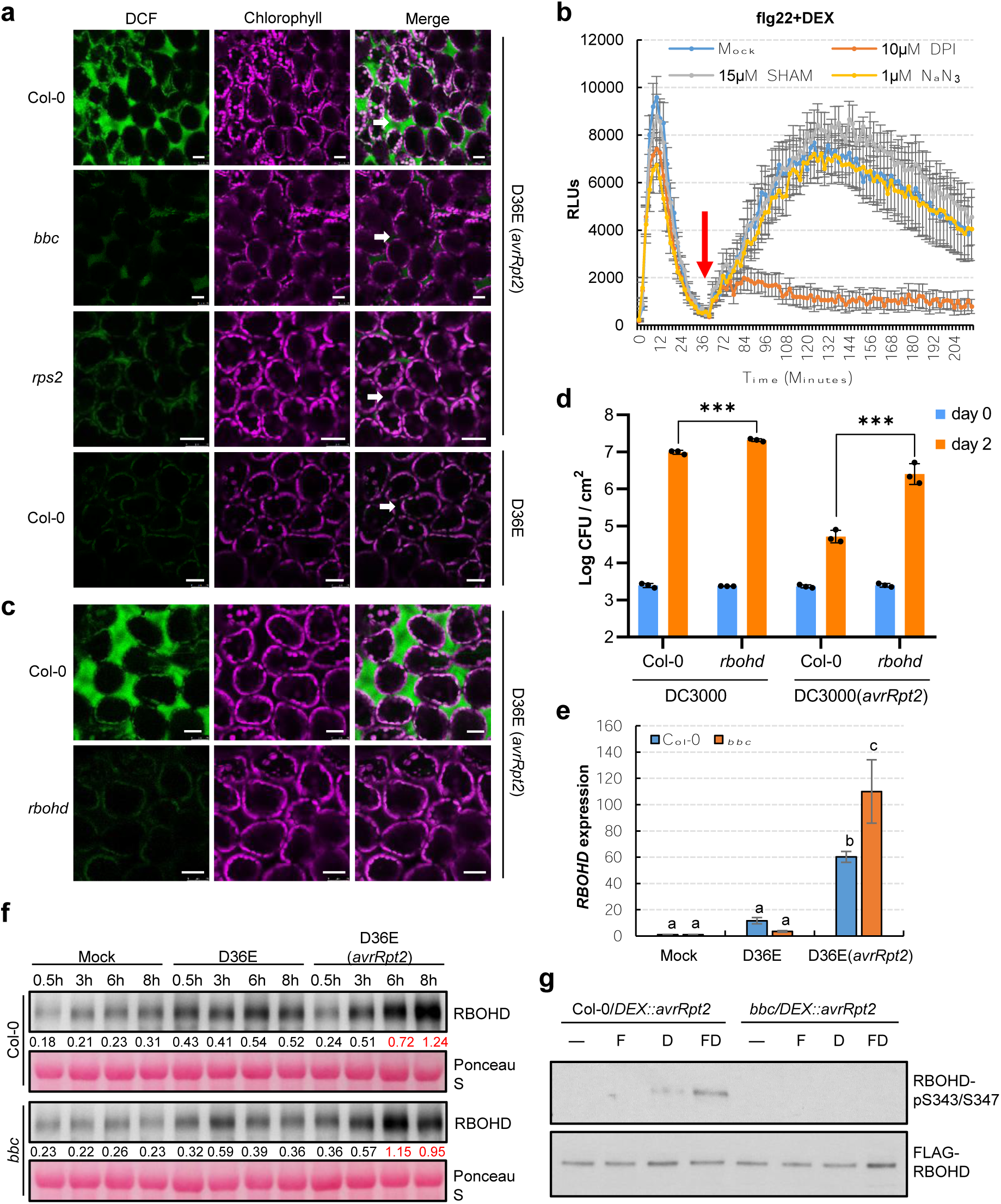
AvrRpt2-triggered ROS is mediated by RBOHD and requires PRR/co-receptors for activation. **a**, ROS burst detected with fluorescent dye H_2_DCFDA in Col-0, *bbc, rps2* and *rbohd* leaves 5h after infiltration of D36E(*avrRpt2*) or in Col-0 leaves 5h after infiltration of D36E strain. White arrows indicate the apoplast space in the leaf. Scale bars = 25μm. **b**, ETI-associated ROS burst is inhibited by DPI, an NADPH oxidase inhibitor. ROS was detected in Col-0/*DEX::avrRpt2* plants after treatment of 100nM flg22 and 5μM DEX. Chemical inhibitors (DPI, SHAM or NaN_3_) were added after the first ROS burst (about 40min after addition of flg22 and DEX). Data are displayed as mean ± s.e.m. (*n* ≥ 6). **c**, ROS burst detected with fluorescent dye H_2_DCFDA in Col-0 and *rbohd* leaves 5h after infiltration of D36E(*avrRpt2*). Scale bars = 25μm. **d**, The *rbohd* mutant plant is compromised in ETI resistance against *Pst* DC3000 (*avrRpt2*). Bacteria were infiltrated into Arabidopsis leaves at OD_600_=0.001 and bacterial populations inside leaves were determined 2 days post infection. ***, student’s *t*-test, two-tailed, *p* < 0.001. Data are displayed as mean ± s.d. (*n* = 3). **e**, RBOHD transcript levels in Col-0 and *bbc* plants 3h after inoculation of bacterial strains indicated. Data are displayed by mean ± s.e.m. (*n* ≥ 3). Different letters indicate statistically significant differences, as analyzed by two-way ANOVA (*p* < 0.05). **f**, RBOHD protein levels in Col-0 and *bbc* plants at different time points after inoculation of bacterial strains indicated. The numbers indicate band intensity relative to that of Ponceau S, quantified by ImageJ, and red indicates strong induction. **g**, Phosphorylation of RBOHD protein at S343/S347 sites requires PRR/co-receptors. FLAG-RBOHD was transformed into protoplasts prepared from Col-0/*DEX::avrRpt2* and *bbc*/*DEX::avrRpt2* plants. Protoplasts were treated with elicitors (—, Mock; F, 100nM flg22; D, 5μM DEX; FD, 100nM flg22+5μM DEX) and harvested 2.5h later for FLAG-RBOHD immunoprecipitation. Phosphorylated and total RBOHD proteins were detected by RBOHD pS343/347-specific antibody and FLAG antibody, respectively. All experiments were repeated at least three times with similar trends.

### Coordination of PRR/co-receptors and NLR for activation of RBOHD

We next assayed the transcript and protein level of RBOHD and found that the transcript (Fig. 2e) and protein level (Fig. 2f) of RBOHD are induced both by D36E and, interestingly, to a much higher level, by D36E(*avrRpt2*) inoculation in Col-0 plant. Surprisingly, the strong induction of *RBOHD* transcript and protein by D36E(*avrRpt2*) occurred at a comparable level in *bbc* mutant plants (Fig. 2e, f). These results indicate that neither *RBOHD* transcript nor RBOHD protein accumulation accounts for the compromised ROS production in the *bbc* mutant and suggests an involvement of PRRs/PTI in post-translational regulation of RBOHD during ETI. Previous studies have reported several classes of kinases, including calcium dependent protein kinases (CPKs) and *Botrytis*-induced kinase 1 (BIK1), involved in phosphorylating RBOHD for ROS production^33-37^. We examined the ETI-ROS level in the *cpk5/6/11* quadruple mutant and *bik1* mutant plants, and found that ETI-associated ROS was reduced in *bik1* mutant but did not seem to be affected in *cpk5/6/11* mutant plant (Extended Data Fig. 6), suggesting that BIK1 contributes to the production of ETI-ROS. BIK1 was reported to rapidly and transiently (i.e. at 15min post-elicitation) phosphorylate RBOHD at multiple sites including S39, S343 and S347 during PTI activation^33,34^. We therefore examined RBOHD phosphorylation levels in protoplasts prepared from Col-0/*DEX::avrRpt2* and *bbc*/*DEX::avrRpt2* plants and transformed with an DNA construct expressing FLAG-RBOHD. PTI and ETI in these protoplasts were activated by flg22 and DEX, respectively. A 35S promoter was used to express FLAG-RBOHD to ensure similar protein levels during various treatments. While no phosphorylation at S343/S347 in Col-0 protoplasts was detected 2.5h after treatment with flg22, DEX alone reproducibly induced a weak phosphorylation of S343/S347 in Col-0 protoplasts 2.5h after treatment (Fig. 2g). Strikingly, a flg22+DEX treatment induced a much stronger phosphorylation on S343/S347 in Col-0 background 2.5h after treatment (Fig. 2g), suggesting a synergistic effect of PTI and ETI in phosphorylating RBOHD at S343/S347. In contrast, no phosphorylation was detected in the *bbc* background with flg22, DEX or flg22+DEX treatment, confirming the requirement of PRR/co-receptors for RBOHD phosphorylation during ETI. Our results illustrate the importance of two classes of immune receptors in the coordination of the abundance (i.e. by RPS2) and full activity (i.e. by PRRs) of RBOHD for generating robust ETI-ROS. Interestingly, S343/S347 phosphorylation of RBOHD has previously been shown to be important for ETI resistance and restriction of bacterial growth^38^.

### Examination of PTI- and ETI-associated transcriptome

The requirement of PRR/co-receptors for activation of RBOHD and a strong up-regulation of RBOHD during ETI (Fig. 2e, f) were intriguing to us and suggested that ETI may have evolved to co-opt RBOHD and other components of the PTI pathway as an integral part of its signaling mechanism. We therefore examined the expression patterns of other components of the PTI pathway and the rest of Arabidopsis transcriptome by RNAseq (Fig. 3a). Bacteria were infiltrated at a high dose (i.e., ∼2×10^7^ cfu/mL) and expression was examined at early time points (i.e. 3h and 6h post infiltration) to ensure similar bacterial populations in Col-0 and *bbc* plants at the sampling times (Extended Data Fig. 7a). We found that, at 3h post infiltration, D36E(*avrRpt2*) already caused significant differential expression for many genes (i.e., more than 4,000 genes) compared to D36E in Col-0 plant (Extended Data Fig. 7b), suggesting that 3h is sufficient for delivery of AvrRpt2 into the plant cell and triggering strong ETI-associated gene expression. We therefore focused analysis on 3h time point in order to reveal early, and likely more direct, changes of ETI-associated gene expression. Many genes are differentially regulated at this early time point between Col-0 and *bbc* plants in response to PTI-inducing D36E (Fig. 3b), as expected. Interestingly, the majority of these genes show similar expression pattern in Col-0 and *bbc* plants after D36E(*avrRpt2*) inoculation (Fig. 3b), suggesting that ETI can largely restore PTI-associated global gene expression in the *bbc* plant. Similar trends were observed for genes associated with salicylic acid, jasmonate and ethylene pathways (Extended Data Fig. 7c-e). We did notice that a subset of 272 genes were differentially expressed in *bbc* plants after D36E(*avrRpt2*) inoculation (Supplementary table 1). In particular, a cluster of *WRKY* genes including *WRKY22/29* and *FRK1* (*Flg22-induced Receptor-like Kinase 1*), which are canonical marker genes of flg22-induced PTI pathway^39^, are down-regulated in the *bbc* plant (Extended Data Fig. 8a, b). This suggests that, despite the general rescue of PTI-associated gene expression by ETI, the WRKY-FRK1 branch represents a unique immune branch, the activation of which during ETI requires PRR/co-receptors.

**Fig.3.**
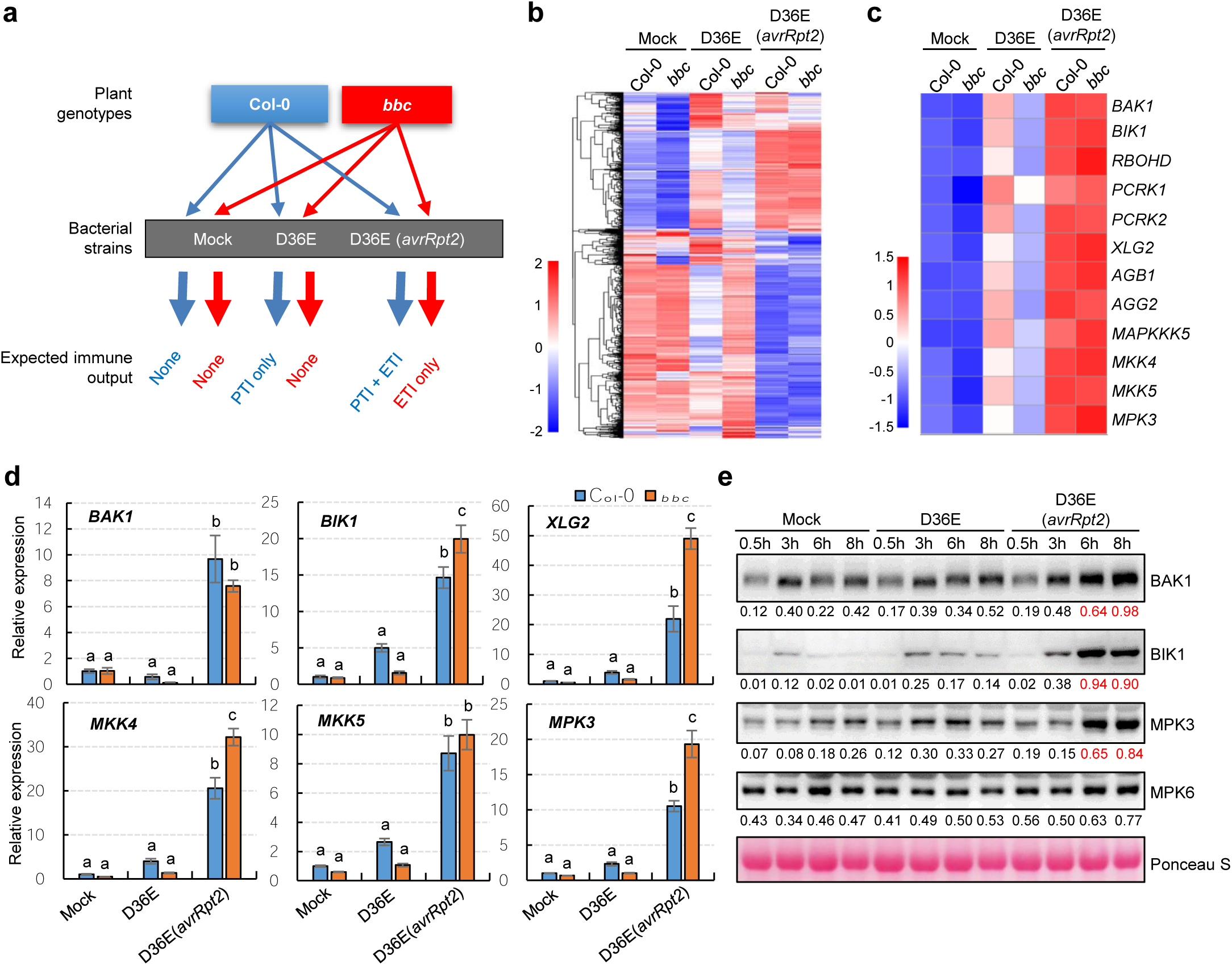
ETI upregulates key components of the PTI pathway. **a**, A diagram showing the RNAseq design in this study. **b**, A heat-map of the expression pattern of D36E/PTI-responsive genes in the RNAseq experiment. **c**, A heat map of the expression pattern of PTI pathway genes, showing restoration and up-regulation of expression of major PTI components during AvrRpt2-triggered ETI. **d**, qRT-PCR results of representative PTI pathway genes. Col-0 and *bbc* plants were infiltrated with different strains indicated, and leaves were harvested 3h post infiltration for transcript analysis (mean ± s.e.m; *n* ≥ 3). Different letters indicate statistically significant differences as analyzed by two-way ANOVA (*p* < 0.05). **e**, Protein levels of BAK1, BIK1, MPK3 and MPK6 in Col-0 plants at different time points after inoculation of bacterial strains indicated. BAK1 is detected in the immunoblot of total membrane fraction and other proteins are detected in the immunoblot of total protein extracts. Equal amounts of total proteins were loaded into gel lanes. MPK6 protein is not induced by ETI and serves as an internal control. The numbers indicate band intensity relative to that of Ponceau S, quantified by ImageJ, and red indicates strong induction. All experiments were repeated at least three times with similar trends.

### Augmentation of key PTI components by ETI

Further analysis of PTI- and ETI-associated transcriptomes revealed an interesting expression pattern for many PTI signaling genes. As shown in Fig. 3c, while PTI-inducing D36E can induce moderately many key PTI components, namely *BAK1, BIK1, XLG2/AGB1/AGG2*^40^, *MKK4/5* and *MPK3*, ETI-inducing D36E(*avrRpt2*) induced these genes to a much higher level. Similar to *RBOHD*, the strong activation of these PTI components by ETI is independent of PRR/co-receptors, since it occurs in the *bbc* mutant. Noticeably, BIK1 and some other PBLs, but not PBL1, are strongly induced after D36E(*avrRpt2*) inoculation (Extended Data Fig. 9), suggesting differential contribution of different members of the BIK1/PBL family to ETI. Quantitative RT-PCR was performed to confirm the RNAseq results (Fig. 3d), and western blot further confirmed, at the protein level, the ETI up-regulation of several key components, including BAK1, BIK1 and MPK3 (Fig. 3e). Our results suggest that part of the AvrRpt2-ETI response is to ensure rapid high-level expression of key components of the PTI pathway, consistent with PTI being an essential component of ETI. We propose that this ETI-mediated up-regulation of PTI components is also likely an important part of a mechanism to overcome the negative regulation of PTI by endogenous “braking” systems of plants and exogenous pathogen effectors during infections^2,41^.

A previous study showed that SA signaling could up-regulate PRR protein level at a later stage (i.e., 24h after benzothiadiazole treatment)^42^. We therefore tested gene expression in the SA-deficient *sid2* plant and found that PTI components were still up-regulated by D36E(*avrRpt2*) (3h after inoculation) (Supplementary Fig. 10a). In addition, we examined our RNAseq dataset for the expression patterns of responsive genes to SA and N-hydroxy-pipecolic acid (NHP), which have been shown to function synergistically in plant immunity^43,44^. Our results showed that both SA-(Extended Data Fig. 7c) and NHP-(Extended Data Fig. 10b) responsive genes had similar transcript levels in Col-0 and *bbc* plants after D36E(*avrRpt2*) inoculation, suggesting intact SA/NHP signaling in the *bbc* mutant during AvrRpt2-ETI. Therefore, ETI appears to rapidly “re-enforce” the PTI pathway in a SA/NHP-independent manner.

## Discussion

Our study reveals a surprising requirement of PRR/co-receptors for effective ETI and supports a mechanistic model in which ETI co-opts the PTI machinery, including the BIK1-RBOHD module, as an indispensable component (Fig. 4). In particular, we found that PRRs and NLRs, the two primary classes of plant immune receptors, function synergistically to ensure a fully “active status” as well as “robust level” of a key immune component, RBOHD, which mediates ETI-ROS generation and full disease resistance. We also identified PRR-mediated RBOHD phosphorylation at S343/S347 sites as one of the mechanistic links between PTI and ETI.

**Fig.4.**
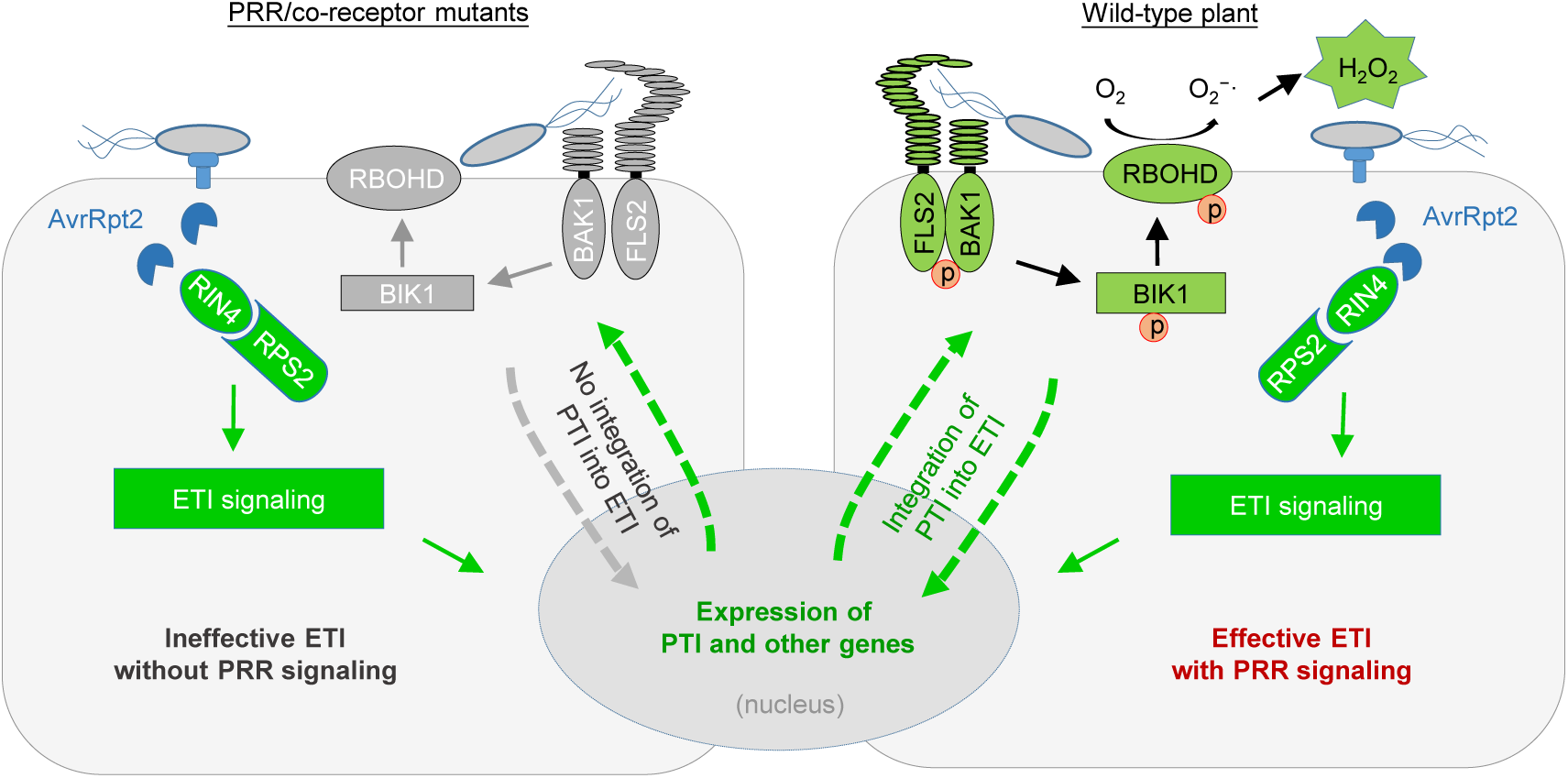
A model depicting findings from this study showing PTI as a key component of ETI. Grey color indicates mutated (i.e. FLS2 and BAK1) or inactive (i.e. RBOHD and BIK1) proteins. In wild-type plant, PTI is integrated into ETI in that RPS2 activation leads to protein accumulation of PTI components such as RBOHD, and PRR/co-receptors are required for fully “activating” it by phosphorylation. In the absence of PRR/co-receptors (left panel), although NLR activation strongly induces protein accumulation of PTI components, many of these components, such as BIK1 and RBOHD, are inactive (shown by the grey arrows), leading to compromised ETI resistance.

Our study sheds light on a long-standing puzzle in the field of plant immunity with respect to the enigmatic similarities in many PTI- and ETI-associated cellular defense features. Our model is supported by a parallel study by Ngou et al. (see back-to-back submission), who focus on the *P. syringae* effector AvrRps4 recognized by TIR-type NLRs (RPS4/RRS1), while we focus on RPS2, a CC-type NLR. Our complementary data suggest conservation of the discovered mechanism for two different types of NLRs, which account for the vast majority of pathogen-sensing NLRs in the plant kingdom. Intriguingly, synergistic interaction between cell surface and intracellular immune receptors in animals and humans has also been reported^45-48^, suggesting a possible conceptual parallel in immune receptor functions across kingdoms of life.

The demonstration of PTI as an integral component of ETI has significant implications in understanding how ETI resistance mechanisms prevent pathogen growth. Specifically, several PTI-associated anti-pathogen mechanisms have been described recently, including suppression of bacterial type III secretion^49-51^, inhibition of bacterial motility by lignification^52^ and restriction of nutrient acquisition^53^. Our study suggests that ETI likely co-opts these PTI anti-pathogen mechanisms to halt pathogen growth. Our work could also have broad practical implications as well, as it suggests a possibility for carefully controlled augmentation of PTI components as a new strategy to broadly increase the effectiveness of ETI against numerous diseases in crop plants.

## Methods

### Plant materials and growth conditions

*Arabidopsis thaliana* plants used in this study are in Col-0 ecotype background. The *fls2/efr/cerk1*^54^, *bak1/bkk1/cerk1*^16^, *rps2*^55^, *rbohd*^30^, *bik1*^56^, *cpk5/6/11*^36^ mutants were reported previously. Plants were grown in potting soil in environmentally-controlled growth chambers, with relative humidity set at 60% and temperature at 22°C with a 12h light/12h dark photoperiod unless stated otherwise. Four-to five-week-old plants were used for all experiments in this study. To generate the *bbc*/*DEX::avrRpt2* and Col-0/*DEX::avrRpt2* transgenic plants, the *avrRpt2* gene was cloned into pBUD-DEX (pBD) vector in the *Xho*I/*Spe*I restriction enzyme sites, and the expression cassette was introduced into Col-0 or *bbc* plants by *Agrobacterium*-mediated transformation.

### Bacterial disease and HR assays

The *Pst* DC3000 strains carrying *avrRpt2, avrRps4* and *avrPphB* were published previously^57-59^. The D36E*(avrRpt2)* strain was generated by transforming the *avrRpt2* expression plasmid into D36E strain by electroporation. For bacterial inoculation, *Pst* strains were cultured in Luria-Marine (LM) medium overnight at 30°C to a cell density of OD_600_=0.8-1.0. Bacteria were collected by centrifugation and washed once with sterile water, and adjusted to a cell density of OD_600_=0.2. For disease assay, bacterial suspension was further diluted to a cell density of OD_600_=0.001-0.002. Bacteria were infiltrated into leaves with a needleless syringe, and inoculated plants were kept under ambient humidity for about 1h to allow evaporation of excess water from the leaf and then covered with a transparent plastic dome to keep high humidity for disease to develop. For quantification of bacteria, four leaf discs from two different leaves (after surface sterilization) were taken using a cork borer (7.5mm in diameter) as one biological repeat, and 3-4 repeats were taken for each treatment. Leaf discs were ground and diluted in sterile water, and the extraction solutions were then plated on LM agar plates supplemented with rifampicin (at 50mg/L). Colonies were counted with a stereoscope 24h after incubation at 30°C. For HR assay, *Pst* DC3000(*avrRpt2*) suspension was prepared as described above and bacterial suspension at the cell density of OD_600_=0.2 was syringe-infiltrated into leaves. Plants were then kept under ambient humidity for about 7h before tissue collapse was recorded.

### RIN4 cleavage assays

Arabidopsis plant leaves were infiltrated with *Pst* D36E(*avrRpt2*) (at OD_600_=0.1), and samples were collected at 0, 2, 4, 8h after infiltration by snap-freezing in liquid nitrogen. Three leaves were collected as one biological repeat. Total proteins were extracted in protein extraction buffer (50mM Tris-HCl pH 7.5, 150mM NaCl, 5mM EDTA pH 7.5, 1mM DTT, 1% Triton X-100, 1mM Phenylmethylsulfonyl fluoride) supplemented with 1 x plant protease inhibitor cocktail (Complete EDTA-free, Roche). Cell lysates were centrifuged at 12,000 x *g* for 15min at 4°C, and the pellet was discarded. Protein concentration of the supernatant (“total protein extract”) was determined by Bradford protein assay kit (Bio-Rad). An equal amount of total protein was loaded on 12% SDS acrylamide gels (Bio-Rad) for SDS-PAGE. RIN4 protein was detected by anti-RIN4 antibody at a dilution of 1:1000^60^. Total proteins were stained by Coomassie Brilliant Blue (CBB) to show equal loading.

### MAPK kinase activity assay

Four-week-old plant leaves were infiltrated with *Pst* D36E(*avrRpt2*) (at OD_600_=0.02), and leaves were collected at different time points by snap-freezing in liquid nitrogen. Proteins were extracted in protein extraction buffer (50mM Tris-HCl pH 7.5, 150mM NaCl, 5mM EDTA pH 7.5, 1mM DTT, 1% Triton X-100, 1mM Phenylmethylsulfonyl fluoride) supplemented with 1 x plant protease inhibitor cocktail (Complete EDTA-free, Roche) and 1 x phosphatase inhibitor cocktail (PhosSTOP, Roche). Total protein concentration was determined with Bradford protein assay kit (Bio-Rad). An equal amount of protein was loaded onto 12% SDS-PAGE gel for western blot. Phosphorylated MPK3 and MPK6 proteins were detected by anti-Phospho-p44/42 antibody (Cell Signaling Technology).

### Protein extraction and immunoblotting for PTI signaling components

Four-week-old plant leaves were infiltrated with sterile water (mock) or different *Pst* strains at OD_600_=0.02, and samples were collected at 0.5, 3, 6, 8h after infiltration. Three to four leaves from different plants were collected as one sample. Protein was extracted using Plasma Membrane Protein Isolation Kit (Invent) according to the manufacturer’s protocol. Concentration of the cytosolic protein was determined with Bradford protein assay kit (Bio-Rad). An equal amount of protein was loaded onto SDS-PAGE gel for western blot. Different PTI components were detected by following antibodies with indicated dilution: anti-RBOHD (Agrisera), 1:1000; anti-BAK1 (Agrisera), 1:5000; anti-BIK1 (Agrisera), 1:3000; anti-MPK3 (Sigma-Aldrich), 1:2500; anti-MPK6 (Sigma-Aldrich), 1:5000.

### Protoplast transformation and detection of RBOHD phosphorylation

Protoplasts were prepared from Col-0/*DEX::avrRpt2* and *bbc/DEX::avrRpt2* plants (4-5 weeks old; grown under 10h light/14h dark photoperiod) and transfected with FLAG-RBOHD plasmid. After overnight incubation to allow protein accumulation, protoplasts were treated with 100nM flg22, 5μM DEX or 100nM flg22+5μM DEX and incubated for 2.5h. Total protein was extracted with protein extraction buffer (50 mM HEPES [pH 7.5], 150 mM KCl, 1 mM EDTA, 0.5% Trition-X100, 1 mM DTT, protease inhibitor cocktail), and then incubated with 50μL anti-FLAG M2 agarose beads (Sigma-Aldrich) for 2 h at 4°C. The bound protein was eluted with 50μL of 0.5mg/mL 3xFLAG peptide for 30 min. RBOHD phosphorylation was detected by immunoblotting with RBOHD-pS343/347 antibody published previously^34^.

### ROS detection Assays

ROS measurement with luminol-based approach was performed as previously described with minor modification^33^. Briefly, leaf discs of four-week-old *Arabidopsis* plants were harvested using a cork borer (5.5mm in diameter) and floated on 200μL sterile water in a 96-well plate, and then incubated overnight at room temperature under continuous light. On the next day, water was replaced with a solution containing 30mg/L (w/v) luminol (Sigma-Aldrich) and 20mg/L (w/v) peroxidase from horseradish (Sigma-Aldrich) with 100nM flg22 only, 5μM DEX only or 100nM flg22+5μM DEX. The luminescence was detected for 5-6h with a signal integration time of 2min using Varioskan Flash plate reader (Thermo Fisher Scientific). For determining the effects of chemical inhibitors, 10μM diphenyleneiodonium (DPI; Sigma-Aldrich), 15μM salicylhydroxamic acid (SHAM; Sigma-Aldrich) or 1μM sodium azide was added to the elicitation solution at indicated time points and luminescence was recorded as described above.

For detection of ROS production by 2’,7’-Dichlorofluorescein diacetate (H_2_DCFDA) under confocal microscopy, plants were infiltrated with *Pst* D36E (OD_600_=0.02) or D36E(*avrRpt2*) (OD_600_=0.02), air-dried and put back into the plant growth room. ROS was detected at 4-5h post infiltration. Ten μM of H_2_DCFDA solution was infiltrated into the leaf and fluorescence signal was detected 10 min later. Images were captured using a Leica SP8 microscope with a 488nm excitation and 501-550nm emission, and chlorophyll auto-fluorescence was detected at 640-735nm.

### RNA extraction and qRT-PCR analysis of gene expression

To analyze gene expression levels, four-week-old *Arabidopsis* plant leaves were infiltrated with sterile water (mock) or different *Pst* strains at OD_600_=0.04, and then harvested at indicated time points. Three leaves from different plants were collected as one biological replicate and 4 replicates were collected for each treatment. Samples were frozen and ground in liquid nitrogen. Total mRNA was extracted using Trizol reagent (Invitrogen) according to the manufacturer’s protocol. One μg of RNA was used for reverse transcription using the ReverTra Ace® qPCR RT Master Mix with gDNA remover (TOYOBO). Real-time qPCR analysis was carried out with the SYBR®Green Realtime PCR Master Mix (TOYOBO) on a CFX real-time machine (Bio-Rad). Two technical repeats were performed for each sample. The plant *U-box* gene was used as reference gene for normalization. Primer sequences for qPCR are listed in Supplementary Table 2.

### cDNA library generation and RNAseq

For RNAseq experiments, bacterial inoculation and sample collection were performed as described above. Two leaves from different plants were harvested as one replicate, and four biological replicates were collected for each treatment/time point. Total mRNA was extracted using Trizol reagent (Invitrogen). Total RNA was then treated with DNase I (Invitrogen) to remove DNA and purified RNA was recovered with RNeasy® MinElute™ Cleanup kit (QIAGEN) according to the manufacturer’s instructions. Library construction and RNA sequencing were performed by Novogene company. Briefly, RNA purity and integrity was examined using the NanoPhotometer® spectrophotometer (IMPLEN) and the RNA Nano 6000 Assay Kit of the Bioanalyzer 2100 system (Agilent Technologies). RNA concentration was measured with Qubit® RNA Assay Kit in Qubit® 2.0 Flurometer (Life Technologies). One μg RNA per sample was used as input material for library preparation and sequencing. Sequencing libraries were generated using NEBNext® Ultra™ RNA Library Prep Kit for Illumina® (NEB), following the manufacturer’s recommendations and sequenced on Illumina Hiseq platform and 150 bp paired-end reads were generated.

### Data analysis of RNA-seq

Clean raw data were obtained by removing reads containing adapter sequences or ploy-N and low-quality reads and were then mapped to the Arabidopsis genome (TAIR10). Gene expression levels were calculated using the TPM method (Transcripts per Kb of exon model per Million mapped reads). Differential expression analysis was performed using the DEGSeq R package (1.18.0). The resulting P-values were adjusted using the Benjamini and Hochberg’s approach for controlling the false discovery rate. Genes with an q-value < 0.05 and log_2_(Fold change) > 1 found by DESeq were assigned as differentially expressed.

### Statistical analysis

All statistical analyses were performed by one-way or two-way analysis of variance (ANOVA) with GraphPad software or by two sided student’s *t*-test with Office Excel software. Each experiment was repeated at least three times and data were represented as the mean ± standard error of mean (s.e.m.) or standard deviation (s.d.) as indicated.

### Data availability

All data is available in the main text or the supplementary materials.

## Acknowledgments

We would like to thank Xin lab members for helpful discussions. We thank the Greenhouse and Confocal Microscopy Imaging facilities at CAS Center for Excellence in Molecular Plant Sciences for plant growth and technical support. Dr. Gitta Coaker from University of California, Davis, USA kindly provided RIN4 antibody. We thank Bruno Pok Man Ngou and Pingtao Ding from Jonathan Jones’ lab at the Sainsbury Laboratory, UK, for insightful discussions during manuscript preparation. This research was supported by Chinese Academy of Sciences, Center for Excellence in Molecular Plant Sciences/Institute of Plant Physiology and Ecology, National Key Laboratory of Molecular Plant Genetics and Chinese Academy of Sciences Strategic Priority Research Program (Type-B; Project number: XDB27040211). Guozhi Bi was supported by the Youth Program of National Natural Science Foundation of China (NSFC) (Project number: 31900222).

## Author contributions

With initial observation of PRR dependency for ETI resistance made by X-F. X. while at Michigan State University, supported by the US National Institute of General Medical Sciences (GM109928), M. Y. and X-F. X. designed subsequent experiments at CAS Center for Excellence in Molecular Plant Sciences/Institute of Plant Physiology and Ecology; M. Y., Z. J., G. B., K. N. and M. L. performed all the experiments described; M. Y. and X-F. X. wrote the paper and S. Y. H. and J-M. Z. edited the paper.

## Materials & correspondence

Correspondence and material requests should be addressed to.

---

**Supplementary Table 2.**
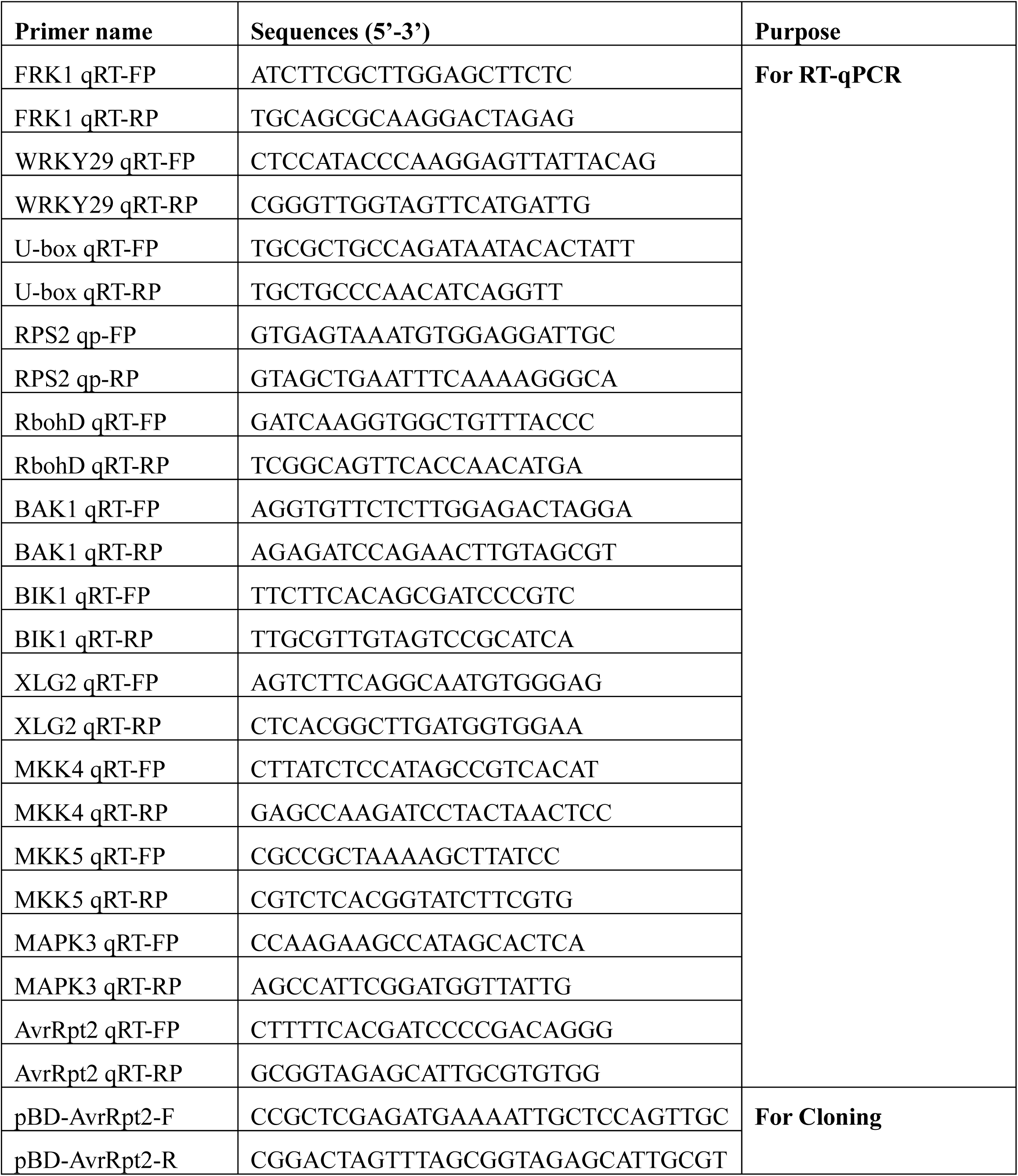
Primers used in this study.

**Extended Data Fig.1.**
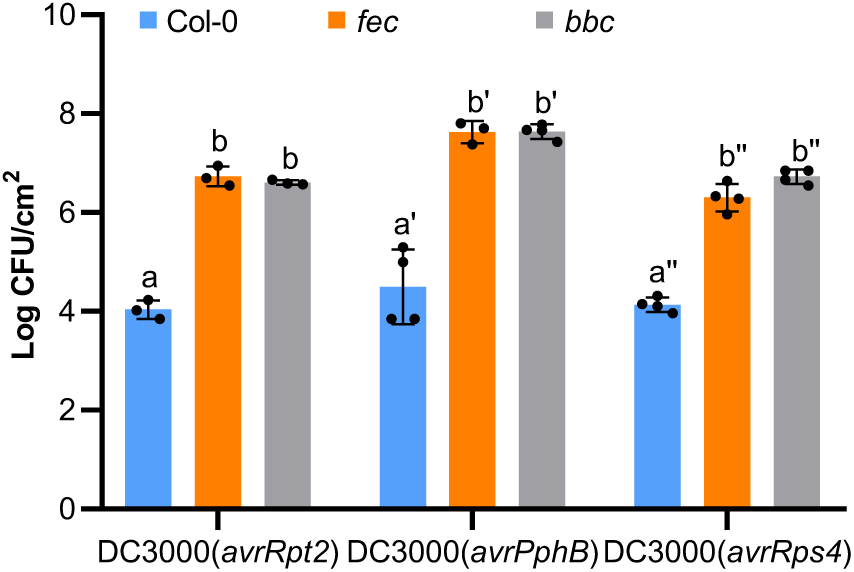
PRR/co-receptors are required for ETI elicited by different *P. syringae* avirulent effectors. AvrPphB- and AvrRps4-mediated ETI are also compromised in *fec* and *bbc* mutants. Plants were infiltrated with different strains at OD_600_=0.002. Bacterial populations were determined 3 days post inoculation. Different letters indicate statistically significant differences in bacterial populations as determined by two-way ANOVA. (mean ± s.d.; *n* ≥ 3; *p* < 0.05). Experiments were repeated at least three times with similar trends.

**Extended Data Fig.2.**
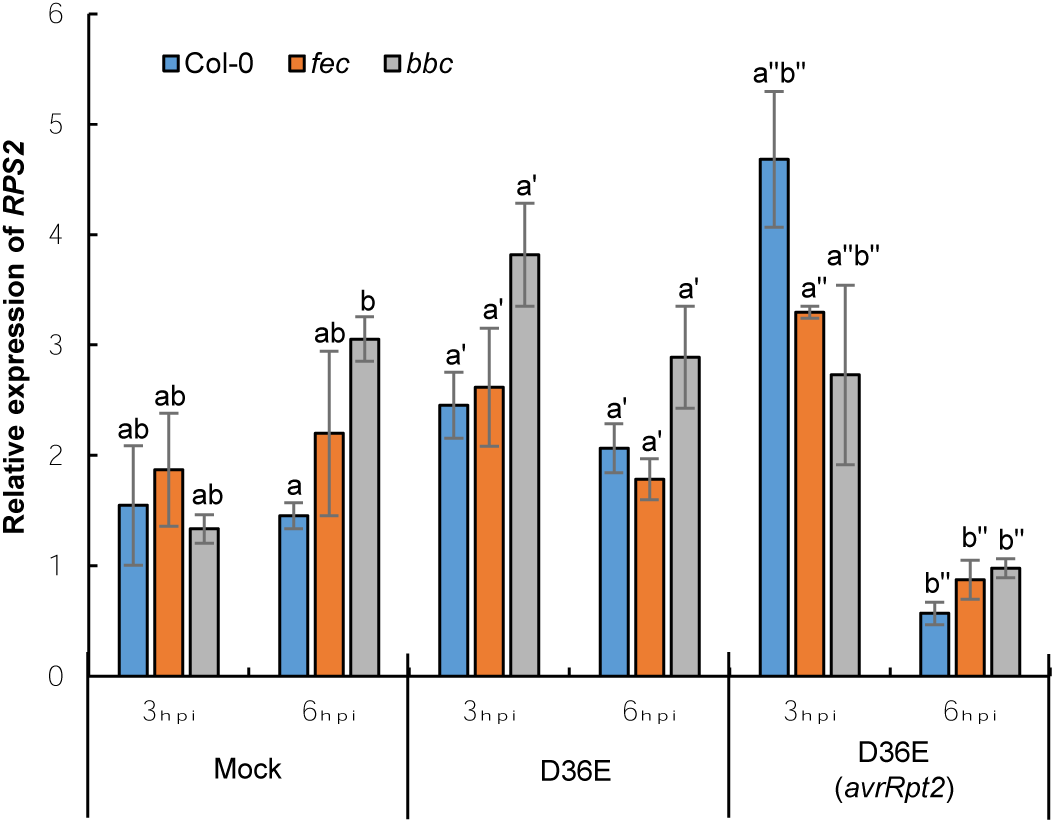
The transcript levels of *RPS2* are not altered in the *fec* and *bbc* mutant plants. *RPS2* transcript levels in the *fec* and *bbc* mutant plants were similar to those in Col-0 plants after inoculation of bacterial strains indicated. Different letters indicate statistically significant differences as analyzed by two-way ANOVA (mean ± s.e.m; *n* = 3; *p* < 0.05). Experiments were repeated at least three times with similar trends.

**Extended Data Fig.3.**
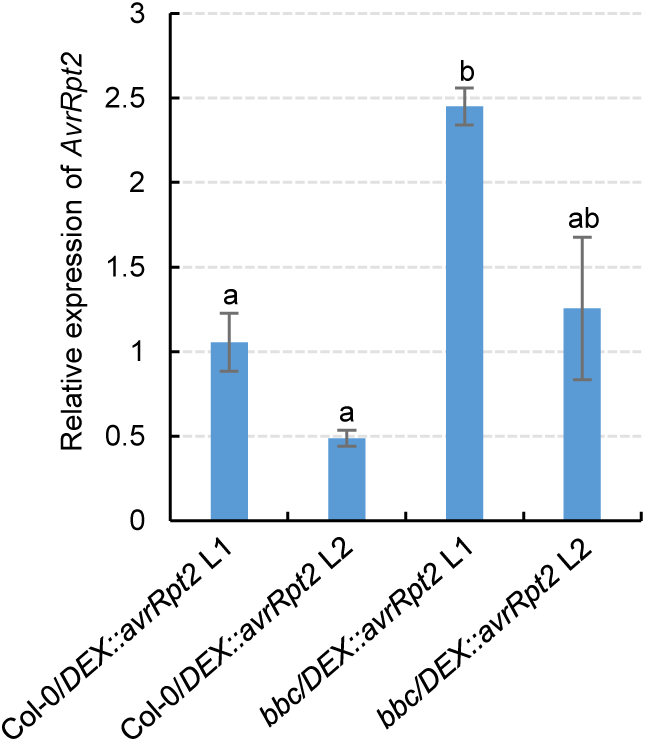
Characterization of different lines of *bbc*/*DEX::avrRpt2* plants. Expression levels of the *avrRpt2* transgene in different transgenic lines 2h after infiltration with 5μM DEX. Different letters indicate statistically significant differences as analyzed by one-way ANOVA (mean ± s.e.m.; *n* ≥ 3; *p* < 0.05). Experiments were repeated at least three times with similar trends.

**Extended Data Fig.4.**
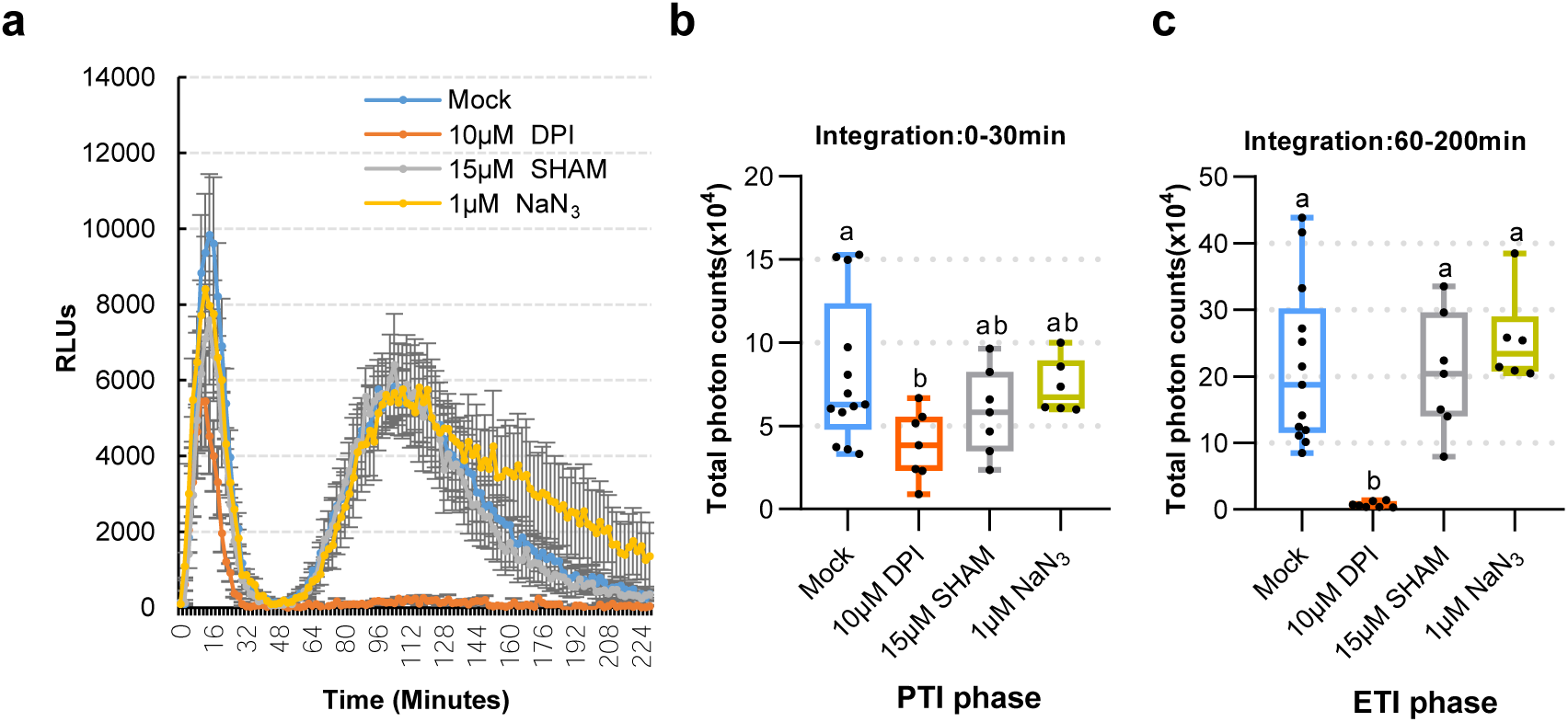
AvrRpt2-triggered ETI-ROS depends on NADPH oxidase. **a-c**, ROS production in Col-0/*DEX::avrRpt2* L1 plants was inhibited by NADPH oxidase inhibitor DPI. Leaf discs were treated with 100nM flg22 and 5μM DEX. DPI, SHAM and NaN_3_ were added at the beginning of measurement (mean ± s.e.m.; *n* ≥ 6). **b-c**, Total photon counts are calculated from **a** at the PTI phase (0-30min) or ETI phase (60-200min). Different letters indicate statistically significant difference as analyzed by one-way ANOVA (*p* < 0.05). Box plots: centre line, median; box limits, lower and upper quartiles; dots, individual data points; whiskers, highest and lowest data points. Experiments were repeated at least three times with similar trends.

**Extended Data Fig.5.**
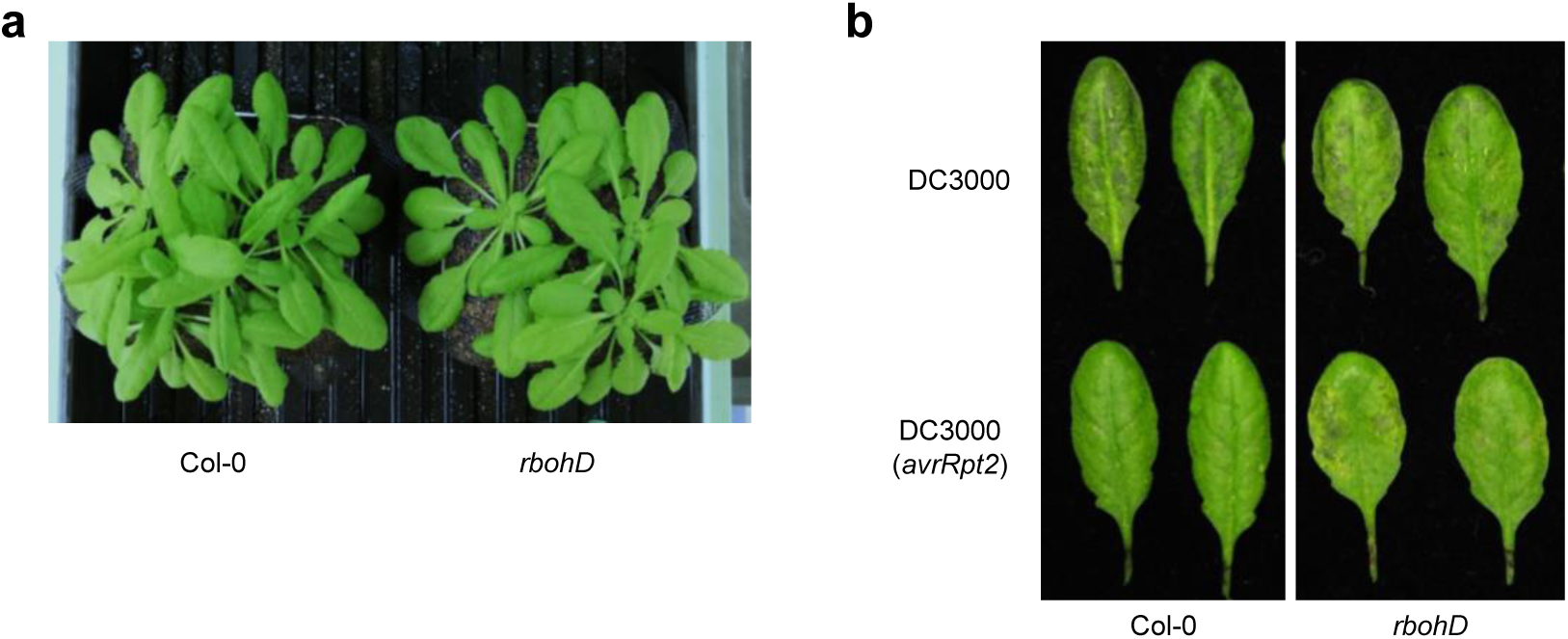
The *rbohd* mutant plant is compromised in ETI resistance against *Pst* DC3000(*avrRpt2*). **a**, Appearance of the 5 week-old *rbohd* mutant plants before bacteria inoculation. **b**, Disease symptom of Col-0 and *rbohd* mutant plant 2 days after *Pst* DC3000 and *Pst* DC3000 (*avrRpt2*) infiltration. Experiments were repeated at least three times with similar trends.

**Extended Data Fig.6.**
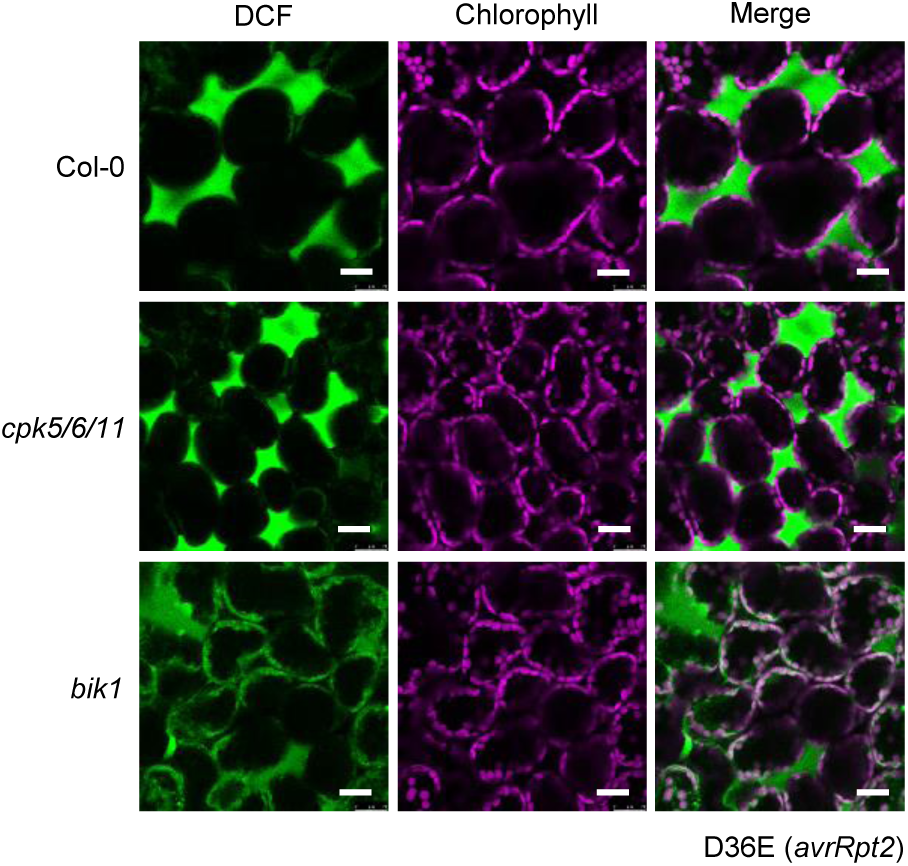
The AvrRpt2 ETI-associated ROS burst is partially mediated by BIK1. ROS was detected in the *bik1* and *cpk5/6/11* mutant plants by H_2_DCFDA dye 4.5 h after D36E (*avrRpt2*) inoculation. Scale bars = 25 μm. Experiments were repeated at least three times with similar trends.

**Extended Data Fig.7.**
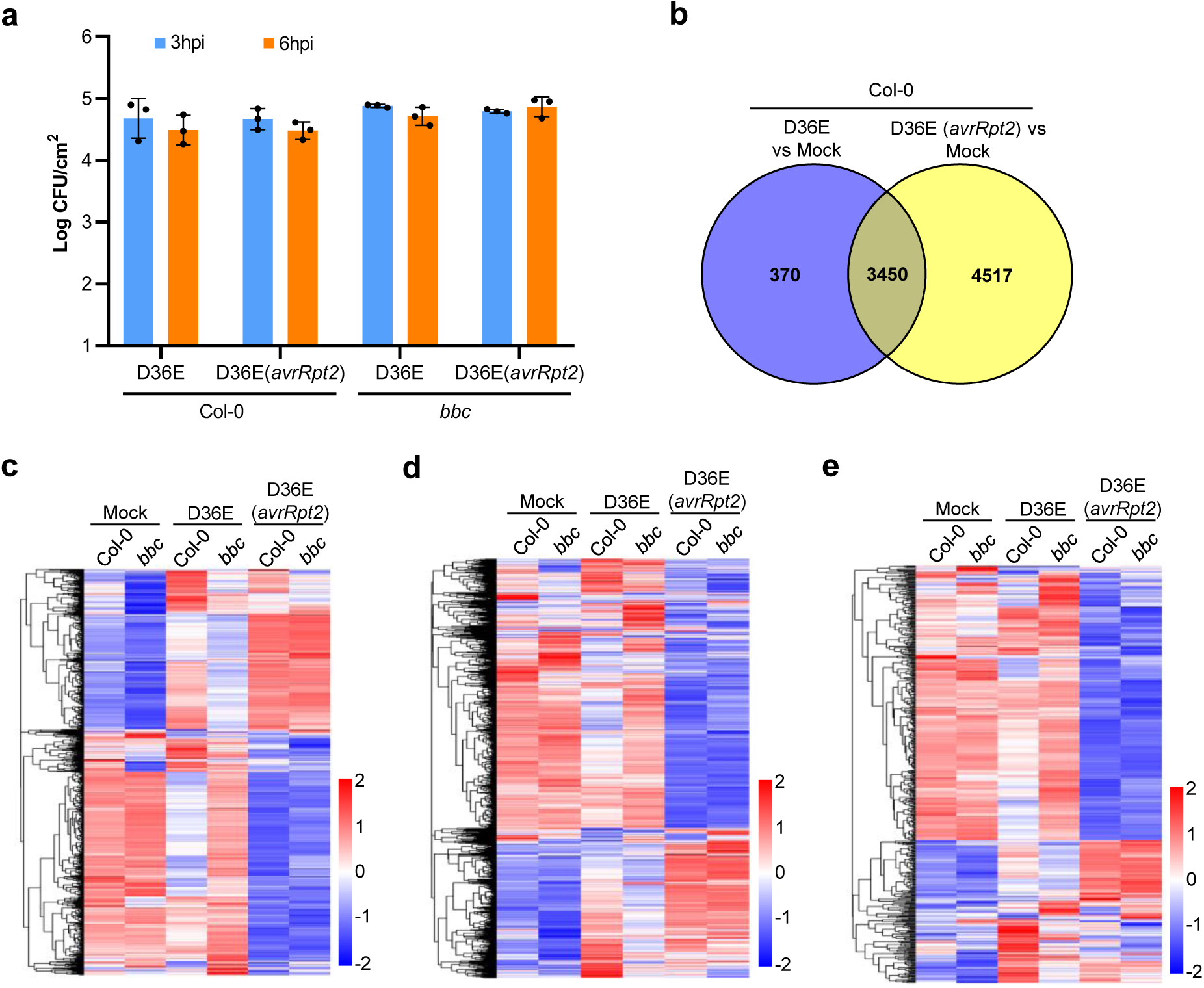
Transcriptomic analysis of RNAseq experiments. **a**, Bacterial population in Arabidopsis leaves at 3h or 6h post infiltration. Data are displayed by mean ± s.d. (n = 3). **b**, A Venn diagram showing numbers of differentially expressed genes (DEGs) 3h after D36E or D36E(*avrRpt2*) infection in Col-0 plants. **c-e**, Heat-maps of SA-**(c**; genes extracted from Karolina et al., 2012^61^), jasmonate-**(d;** genes extracted from Hickman et al., 2017^62^) and ethylene-**(e;** genes extracted from Nemhauser et al.,2006^63^) responsive genes.

**Extended Data Fig.8.**
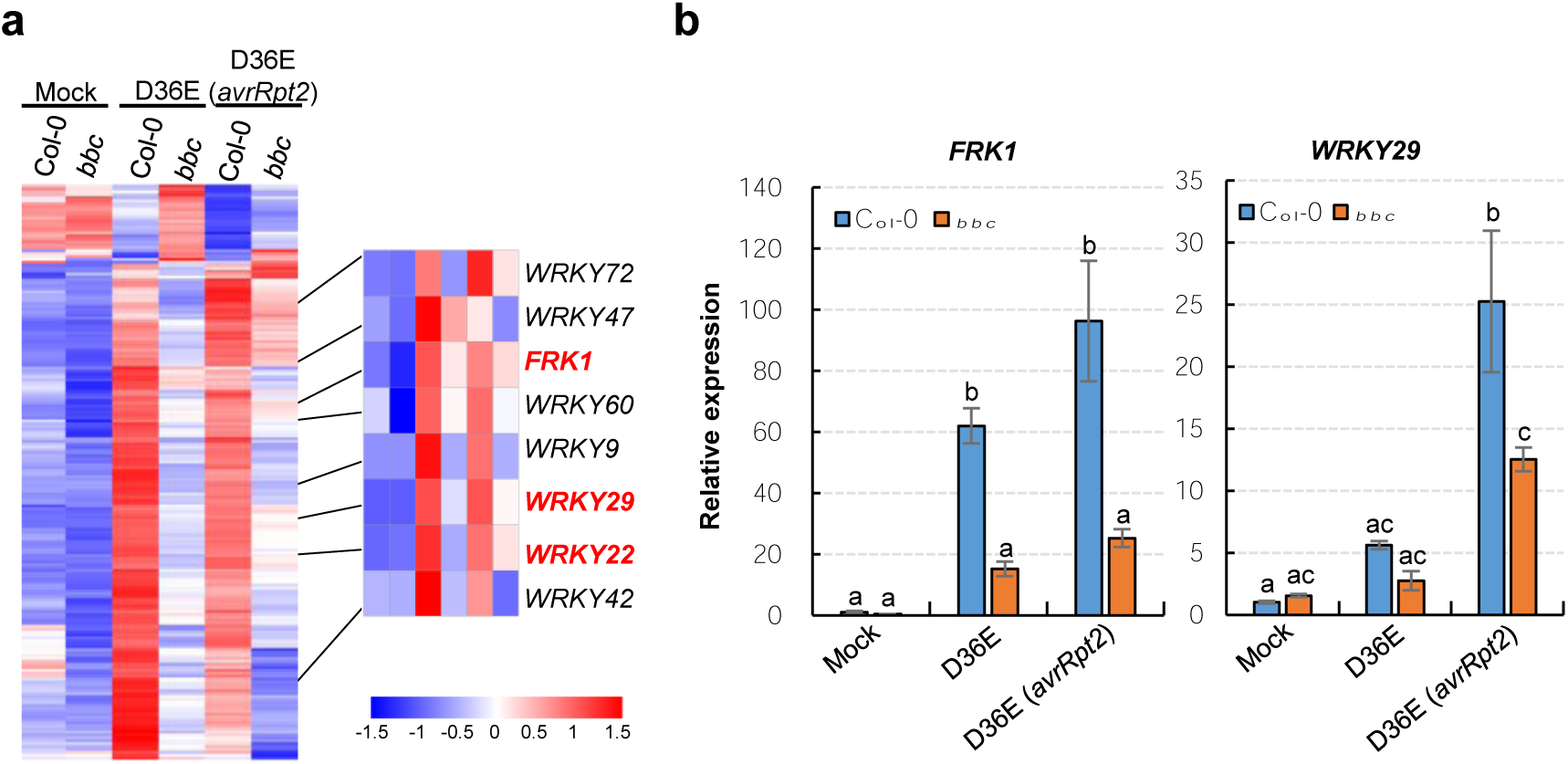
The WRKY-FRK1 is a unique immune branch and can not be restored by ETI in *bbc* mutant. **a**, Heat map of the 272 DEGs in the *bbc* plant compared to Col-0 plant after D36E (*avrRpt2*) infection, with the canonical PTI pathway genes highlighted in red. **b**, qRT-PCR of *FRK1* and *WRKY29* expression level in Col-0 and *bbc* plants 3h after infiltration with different strains or Mock. (mean ± s.e.m.; *n* = 3; statistical analysis by two-way ANOVA; *p* < 0.05; different letters indicate statistically significant difference). Experiments were repeated at least three times with similar trends.

**Extended Data Fig.9.**
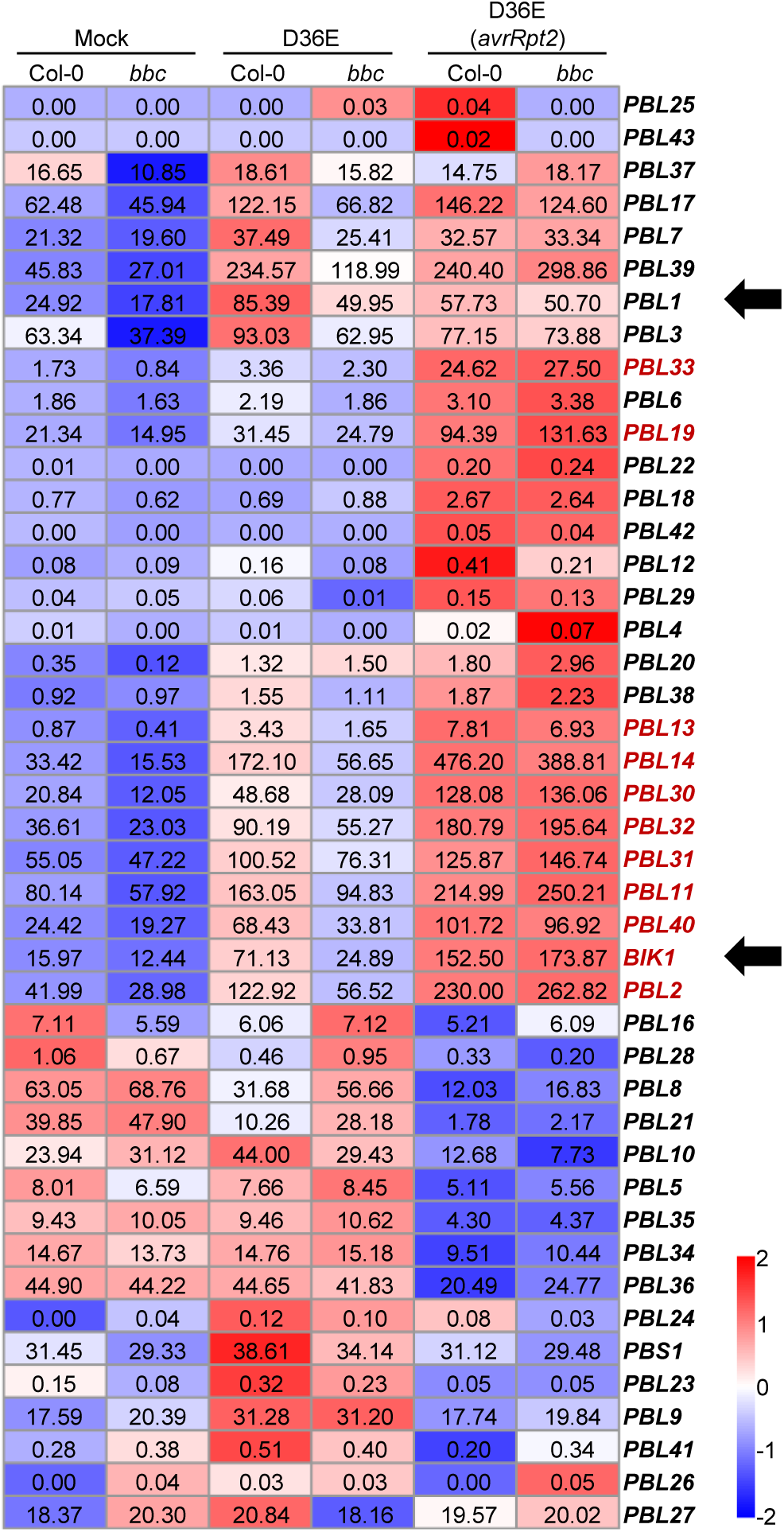
Heat map of *BIK1*/*PBL* family gene expression in the RNAseq experiment. Numerical values indicate expression level calculated by TPM (Transcripts per Kb of exon model per Million). Genes labeled in red show significant up-regulation after D36E(*avrRpt2*) inoculation, compared to mock and D36E inoculation, in Col-0 and *bbc* plants. Arrows indicate *BIK1* and *PBL1* genes.

**Extended Data Fig.10.**
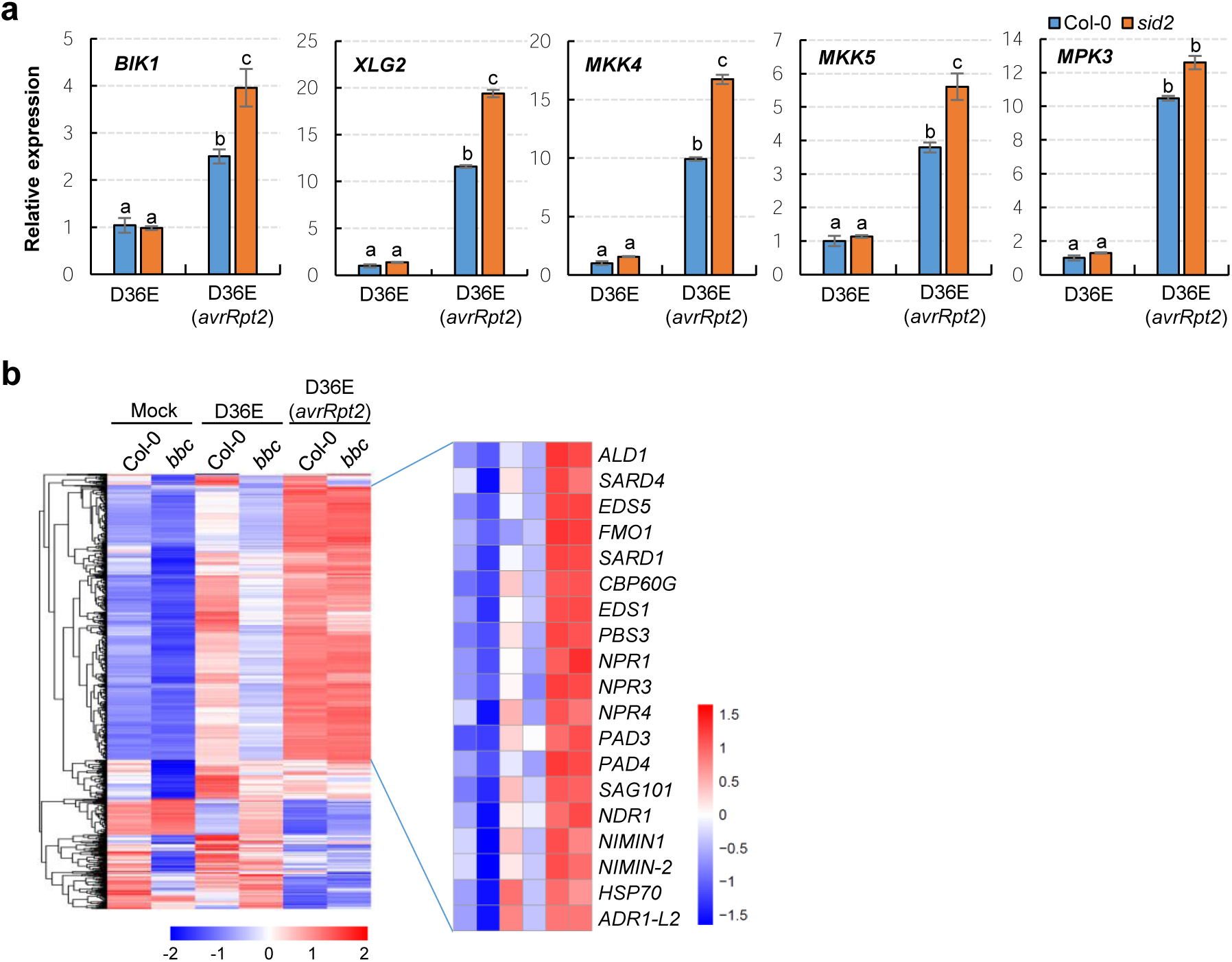
Up-regulation of key PTI component genes by AvrRpt2-triggered ETI seems to be independent of SA/NHP. **a**, qRT-PCR analysis of *BIK1, XLG2, MKK4, MKK5* and *MPK3* expression levels in Col-0 and *sid2* plants 3h after infiltration with D36E or D36E(*avrRpt2*). Different letters indicate statistically significant differences, as analyzed by two-way ANOVA (mean ± s.e.m.; n ≥ 3; p < 0.05). Experiments were repeated at least three times with similar trends. **b**, Heat-maps of NHP-responsive genes (extracted from Hartmann et al., 2018^64^, defined by genes that are responsive to pipecolic acid and depend on FMO1 for expression) in the Col-0 and *bbc* plants in our RNAseq experiment.

## References

1 Yu, X., Feng, B., He, P. & Shan, L. From Chaos to Harmony: Responses and Signaling upon Microbial Pattern Recognition. Annu Rev Phytopathol 55, 109–137, doi:10.1146/annurev-phyto-080516-035649 (2017).

2 Couto, D. & Zipfel, C. Regulation of pattern recognition receptor signalling in plants. Nat Rev Immunol 16, 537–552, doi:10.1038/nri.2016.77 (2016).

3 Spoel, S. H. & Dong, X. How do plants achieve immunity? Defence without specialized immune cells. Nat Rev Immunol 12, 89–100, doi:10.1038/nri3141 (2012).

4 Cui, H., Tsuda, K. & Parker, J. E. Effector-triggered immunity: from pathogen perception to robust defense. Annu Rev Plant Biol 66, 487–511, doi:10.1146/annurev-arplant-050213-040012 (2015).

5 Jones, J. D. & Dangl, J. L. The plant immune system. Nature 444, 323–329, doi:10.1038/nature05286 (2006).

6 Chisholm, S. T., Coaker, G., Day, B. & Staskawicz, B. J. Host-microbe interactions: shaping the evolution of the plant immune response. Cell 124, 803–814, doi:10.1016/j.cell.2006.02.008 (2006).

7 Jones, J. D., Vance, R. E. & Dangl, J. L. Intracellular innate immune surveillance devices in plants and animals. Science 354, doi:10.1126/science.aaf6395 (2016).

8 Tsuda, K. & Katagiri, F. Comparing signaling mechanisms engaged in pattern-triggered and effector-triggered immunity. Curr Opin Plant Biol 13, 459–465, doi:10.1016/j.pbi.2010.04.006 (2010).

9 Peng, Y., van Wersch, R. & Zhang, Y. Convergent and Divergent Signaling in PAMP-Triggered Immunity and Effector-Triggered Immunity. Mol Plant Microbe Interact 31, 403–409, doi:10.1094/MPMI-06-17-0145-CR (2018).

10 Wang, J. et al. Reconstitution and structure of a plant NLR resistosome conferring immunity. Science 364, doi:10.1126/science.aav5870 (2019).

11 Wang, J. et al. Ligand-triggered allosteric ADP release primes a plant NLR complex. Science 364, doi:10.1126/science.aav5868 (2019).

12 Wan, L. et al. TIR domains of plant immune receptors are NAD+-cleaving enzymes that promote cell death. Science 365, 799–803, doi:10.1126/science.aax1771 (2019).

13 Horsefield, S. et al. NAD+ cleavage activity by animal and plant TIR domains in cell death pathways. Science 365, 793–799, doi:10.1126/science.aax1911 (2019).

14 Axtell, M. J. & Staskawicz, B. J. Initiation of RPS2-specified disease resistance in Arabidopsis is coupled to the AvrRpt2-directed elimination of RIN4. Cell 112, 369–377, doi:10.1016/s0092-8674(03)00036-9 (2003).

15 Mackey, D., Belkhadir, Y., Alonso, J. M., Ecker, J. R. & Dangl, J. L. Arabidopsis RIN4 is a target of the type III virulence effector AvrRpt2 and modulates RPS2-mediated resistance. Cell 112, 379–389, doi:10.1016/s0092-8674(03)00040-0 (2003).

16 Xin, X. F. et al. Bacteria establish an aqueous living space in plants crucial for virulence. Nature 539, 524–529, doi:10.1038/nature20166 (2016).

17 Shao, F. et al. Cleavage of Arabidopsis PBS1 by a bacterial type III effector. Science 301, 1230–1233, doi:10.1126/science.1085671 (2003).

18 Gassmann, W., Hinsch, M. E. & Staskawicz, B. J. The Arabidopsis RPS4 bacterial-resistance gene is a member of the TIR-NBS-LRR family of disease-resistance genes. Plant J 20, 265–277, doi:10.1046/j.1365-313x.1999.t01-1-00600.x (1999).

19 Xin, X. F., Kvitko, B. & He, S. Y. *Pseudomonas syringae*: what it takes to be a pathogen. Nat Rev Microbiol 16, 316–328, doi:10.1038/nrmicro.2018.17 (2018).

20 Toruno, T. Y., Stergiopoulos, I. & Coaker, G. Plant-Pathogen Effectors: Cellular Probes Interfering with Plant Defenses in Spatial and Temporal Manners. Annu Rev Phytopathol 54, 419–441, doi:10.1146/annurev-phyto-080615-100204 (2016).

21 Roux, M. et al. The Arabidopsis leucine-rich repeat receptor-like kinases BAK1/SERK3 and BKK1/SERK4 are required for innate immunity to hemibiotrophic and biotrophic pathogens. Plant Cell 23, 2440–2455, doi:10.1105/tpc.111.084301 (2011).

22 Wei, H. L. et al. Pseudomonas syringae pv. tomato DC3000 Type III Secretion Effector Polymutants Reveal an Interplay between HopAD1 and AvrPtoB. Cell Host Microbe 17, 752–762, doi:10.1016/j.chom.2015.05.007 (2015).

23 Tsuda, K. et al. Dual regulation of gene expression mediated by extended MAPK activation and salicylic acid contributes to robust innate immunity in Arabidopsis thaliana. PLoS Genet 9, e1004015, doi:10.1371/journal.pgen.1004015 (2013).

24 Qi, J., Wang, J., Gong, Z. & Zhou, J. M. Apoplastic ROS signaling in plant immunity. Curr Opin Plant Biol 38, 92–100, doi:10.1016/j.pbi.2017.04.022 (2017).

25 McNellis, T. W. et al. Glucocorticoid-inducible expression of a bacterial avirulence gene in transgenic Arabidopsis induces hypersensitive cell death. Plant J 14, 247–257, doi:10.1046/j.1365-313x.1998.00106.x (1998).

26 Felix, G., Duran, J. D., Volko, S. & Boller, T. Plants have a sensitive perception system for the most conserved domain of bacterial flagellin. Plant J 18, 265–276, doi:10.1046/j.1365-313x.1999.00265.x (1999).

27 Levine, A., Tenhaken, R., Dixon, R. & Lamb, C. H2O2 from the oxidative burst orchestrates the plant hypersensitive disease resistance response. Cell 79, 583–593, doi:10.1016/0092-8674(94)90544-4 (1994).

28 Chandra, S., Martin, G. B. & Low, P. S. The Pto kinase mediates a signaling pathway leading to the oxidative burst in tomato. Proc Natl Acad Sci U S A 93, 13393–13397, doi:10.1073/pnas.93.23.13393 (1996).

29 Tian, S. et al. Plant Aquaporin AtPIP1;4 Links Apoplastic H2O2 Induction to Disease Immunity Pathways. Plant Physiol 171, 1635–1650, doi:10.1104/pp.15.01237 (2016).

30 Torres, M. A., Dangl, J. L. & Jones, J. D. Arabidopsis gp91phox homologues AtrbohD and AtrbohF are required for accumulation of reactive oxygen intermediates in the plant defense response. Proc Natl Acad Sci U S A 99, 517–522, doi:10.1073/pnas.012452499 (2002).

31 Daudi, A. et al. The apoplastic oxidative burst peroxidase in Arabidopsis is a major component of pattern-triggered immunity. Plant cell 24, 275–287, doi:10.1105/tpc.111.093039 (2012).

32 Li, Y. et al. Glucose triggers stomatal closure mediated by basal signaling through HXK1 and PYR/RCAR receptors in Arabidopsis. J Exp Bot 69, 1471–1484, doi:10.1093/jxb/ery024 (2018).

33 Kadota, Y. et al. Direct regulation of the NADPH oxidase RBOHD by the PRR-associated kinase BIK1 during plant immunity. Mol Cell 54, 43–55, doi:10.1016/j.molcel.2014.02.021 (2014).

34 Li, L. et al. The FLS2-associated kinase BIK1 directly phosphorylates the NADPH oxidase RbohD to control plant immunity. Cell Host Microbe 15, 329–338, doi:10.1016/j.chom.2014.02.009 (2014).

35 Dubiella, U. et al. Calcium-dependent protein kinase/NADPH oxidase activation circuit is required for rapid defense signal propagation. Proc Natl Acad Sci U S A 110, 8744–8749, doi:10.1073/pnas.1221294110 (2013).

36 Gao, X. et al. Bifurcation of Arabidopsis NLR immune signaling via Ca^2+^-dependent protein kinases. PLoS pathog 9, e1003127, doi:10.1371/journal.ppat.1003127 (2013).

37 Zhang, M. et al. The MAP4 Kinase SIK1 Ensures Robust Extracellular ROS Burst and Antibacterial Immunity in Plants. Cell Host Microbe 24, 379–391 e375, doi:10.1016/j.chom.2018.08.007 (2018).

38 Kadota, Y. et al. Quantitative phosphoproteomic analysis reveals common regulatory mechanisms between effector- and PAMP-triggered immunity in plants. New Phytol 221, 2160–2175, doi:10.1111/nph.15523 (2019).

39 Asai, T. et al. MAP kinase signalling cascade in Arabidopsis innate immunity. Nature 415, 977–983, doi:10.1038/415977a (2002).

40 Liang, X. et al. Arabidopsis heterotrimeric G proteins regulate immunity by directly coupling to the FLS2 receptor. Elife 5, e13568, doi:10.7554/eLife.13568 (2016).

41 Macho, A. P. & Zipfel, C. Targeting of plant pattern recognition receptor-triggered immunity by bacterial type-III secretion system effectors. Curr Opin Microbiol 23, 14–22, doi:10.1016/j.mib.2014.10.009 (2015).

42 Tateda, C. et al. Salicylic acid regulates Arabidopsis microbial pattern receptor kinase levels and signaling. Plant cell 26, 4171–4187, doi:10.1105/tpc.114.131938 (2014).

43 Sun, T. et al. Redundant CAMTA Transcription Factors Negatively Regulate the Biosynthesis of Salicylic Acid and N-Hydroxypipecolic Acid by Modulating the Expression of SARD1 and CBP60g. Mol Plant 13, 144–156, doi:10.1016/j.molp.2019.10.016 (2020).

44 Kim, Y., Gilmour, S. J., Chao, L., Park, S. & Thomashow, M. F. Arabidopsis CAMTA Transcription Factors Regulate Pipecolic Acid Biosynthesis and Priming of Immunity Genes. Mol Plant 13, 157–168, doi:10.1016/j.molp.2019.11.001 (2020).

45 Rathinam, V. A. et al. TRIF licenses caspase-11-dependent NLRP3 inflammasome activation by gram-negative bacteria. Cell 150, 606–619, doi:10.1016/j.cell.2012.07.007 (2012).

46 Baroja-Mazo, A. et al. The NLRP3 inflammasome is released as a particulate danger signal that amplifies the inflammatory response. Nat Immunol 15, 738–748, doi:10.1038/ni.2919 (2014).

47 Franklin, B. S. et al. The adaptor ASC has extracellular and ‘prionoid’ activities that propagate inflammation. Nat Immunol 15, 727–737, doi:10.1038/ni.2913 (2014).

48 Cao, X. Self-regulation and cross-regulation of pattern-recognition receptor signalling in health and disease. Nat Rev Immunol 16, 35–50, doi:10.1038/nri.2015.8 (2016).

49 Crabill, E., Joe, A., Block, A., van Rooyen, J. M. & Alfano, J. R. Plant immunity directly or indirectly restricts the injection of type III effectors by the *Pseudomonas syringae* type III secretion system. Plant physiol 154, 233–244, doi:10.1104/pp.110.159723 (2010).

50 Anderson, J. C. et al. Decreased abundance of type III secretion system-inducing signals in Arabidopsis *mkp1* enhances resistance against *Pseudomonas syringae*. Proc Natl Acad Sci U S A 111, 6846–6851, doi:10.1073/pnas.1403248111 (2014).

51 Nobori, T. et al. Transcriptome landscape of a bacterial pathogen under plant immunity. Proc Natl Acad Sci U S A 115, E3055–E3064, doi:10.1073/pnas.1800529115 (2018).

52 Lee, M. H. et al. Lignin-based barrier restricts pathogens to the infection site and confers resistance in plants. EMBO J 38, e101948, doi:10.15252/embj.2019101948 (2019).

53 Yamada, K., Saijo, Y., Nakagami, H. & Takano, Y. Regulation of sugar transporter activity for antibacterial defense in Arabidopsis. Science 354, 1427–1430, doi:10.1126/science.aah5692 (2016).

## References

54 Gimenez-Ibanez, S., Ntoukakis, V. & Rathjen, J. P. The LysM receptor kinase CERK1 mediates bacterial perception in Arabidopsis. Plant Signal Behav 4, 539–541, doi:10.4161/psb.4.6.8697 (2009).

55 Mindrinos, M., Katagiri, F., Yu, G. L. & Ausubel, F. M. The *A. thaliana* disease resistance gene RPS2 encodes a protein containing a nucleotide-binding site and leucine-rich repeats. Cell 78, 1089–1099, doi:10.1016/0092-8674(94)90282-8 (1994).

56 Veronese, P. et al. The membrane-anchored BOTRYTIS-INDUCED KINASE1 plays distinct roles in Arabidopsis resistance to necrotrophic and biotrophic pathogens. Plant cell 18, 257–273, doi:10.1105/tpc.105.035576 (2006).

57 Mudgett, M. B. & Staskawicz, B. J. Characterization of the *Pseudomonas syringae pv. tomato* AvrRpt2 protein: demonstration of secretion and processing during bacterial pathogenesis. Mol Microbiol 32, 927–941, doi:10.1046/j.1365-2958.1999.01403.x (1999).

58 Hinsch, M. & Staskawicz, B. Identification of a new Arabidopsis disease resistance locus, *RPs4*, and cloning of the corresponding avirulence gene, *avrRps4*, from P*seudomonas syringae pv. pisi*. Mol Plant Microbe Interact 9, 55–61, doi:10.1094/mpmi-9-0055 (1996).

59 Aarts, N. et al. Different requirements for EDS1 and NDR1 by disease resistance genes define at least two R gene-mediated signaling pathways in Arabidopsis. Proc Natl Acad Sci U S A 95, 10306–10311, doi:10.1073/pnas.95.17.10306 (1998).

60 Lee, D., Bourdais, G., Yu, G., Robatzek, S. & Coaker, G. Phosphorylation of the Plant Immune Regulator RPM1-INTERACTING PROTEIN4 Enhances Plant Plasma Membrane H^+^-ATPase Activity and Inhibits Flagellin-Triggered Immune Responses in Arabidopsis. Plant cell 27, 2042–2056, doi:10.1105/tpc.114.132308 (2015).

61 Pajerowska-Mukhtar, K. M. et al. The HSF-like transcription factor TBF1 is a major molecular switch for plant growth-to-defense transition. Curr Biol 22, 103–112, doi:10.1016/j.cub.2011.12.015 (2012).

62 Hickman, R. et al. Architecture and Dynamics of the Jasmonic Acid Gene Regulatory Network. Plant cell 29, 2086–2105, doi:10.1105/tpc.16.00958 (2017).

63 Nemhauser, J. L., Hong, F. & Chory, J. Different plant hormones regulate similar processes through largely nonoverlapping transcriptional responses. Cell 126, 467–475, doi:10.1016/j.cell.2006.05.050 (2006).

64 Hartmann, M. et al. Flavin Monooxygenase-Generated N-Hydroxypipecolic Acid Is a Critical Element of Plant Systemic Immunity. Cell 173, 456–469 e416, doi:10.1016/j.cell.2018.02.049 (2018).

